# Uncovering Ghost Introgression Through Genomic Analysis of a Distinct East Asian Hickory Species

**DOI:** 10.1101/2023.06.26.546421

**Authors:** Wei-Ping Zhang, Ya-Mei Ding, Yu Cao, Pan Li, Yang Yang, Xiao-Xu Pang, Wei-Ning Bai, Da-Yong Zhang

**Author notes:** These authors contributed equally to this work (Wei-Ping Zhang, Ya-Mei Ding).

## Abstract

Although the possibility of introgression from ghost lineages (all unsampled extant and extinct taxa) is now widely recognized, detecting and characterizing ghost introgression remains a challenge. Here, we propose a combined use of the popular *D*-statistic method, which tests for the presence of introgression, and the full-likelihood method BPP, which determines which of the possible gene-flow scenarios, including ghost introgression, is truly responsible. We illustrate the utility of this approach by investigating the reticulation and bifurcation history of the genus *Carya* (Juglandaceae), including the beaked hickory *Carya sinensis*. To achieve this goal, we generated two chromosome-level reference genomes respectively for *C. sinensis* and *C. cathayensis*. Furthermore, we re-sequenced the whole genomes of 43 individuals from *C. sinensis* and one individual from each of the 11 diploid species of *Carya*. The latter dataset with one individual per species is used to reconstruct the phylogenetic networks and estimate the divergence time of *Carya*. Our results unambiguously demonstrate the presence of ghost introgression from an extinct lineage into the beaked hickory, dispelling certain misconceptions about the phylogenetic history of *C. sinensis*. We also discuss the profound implications of ghost introgression into *C. sinensis* for the historical biogeography of hickory species. [BPP; *Carya*; *D*-statistic; gene flow; ghost introgression]

## INTRODUCTION

In recent years, increasingly available genome-scale data coupled with advances in analytical methods have revealed that post-speciation gene flow between species, or simply introgression, is ubiquitous across the tree of life (Mallet et al. 2016; Taylor and Larson 2019; Edelman and Mallet 2021). Among many methods for detecting introgression, Patterson’s *D*- statistic (Patterson et al. 2012), also known as ABBA-BABA test (Green et al. 2010; Durand et al. 2011), is perhaps the most popular (Dagilis et al. 2022). The *D*-statistic considers ancestral (A) and derived (B) alleles across the genomes of four taxa: three ingroup taxa and one outgroup, with a ladder-like species tree (Fig. 1). Under the null hypothesis of no gene flow among ingroup taxa, two particular allelic patterns ‘ABBA’ and ‘BABA’ should occur equally frequent (Pamilo and Nei 1988). An excess of either ABBA or BABA, resulting in a *D*-statistic that is significantly different from zero, is usually interpreted as indicative of gene flow between nonsister taxa, i.e., either between P2 and P3 (positive *D*-score) or between P1 and P3 (negative *D*-score) (Fig. 1a). However, an excess of ABBA or BABA can also arise from a ghost lineage introgressing into P1 or P2 (Fig. 1b). More interestingly, Tricou et al.’s (2022a) simulations demonstrate that most significant *D*-statistics are likely attributable to ghost lineages, necessitating a rethink about the usual interpretation of the *D*-statistic. In other words, failure to consider ghost introgression as a plausible working hypothesis will probably lead to errors of inference in any study of introgression.

**FIGURE 1.**
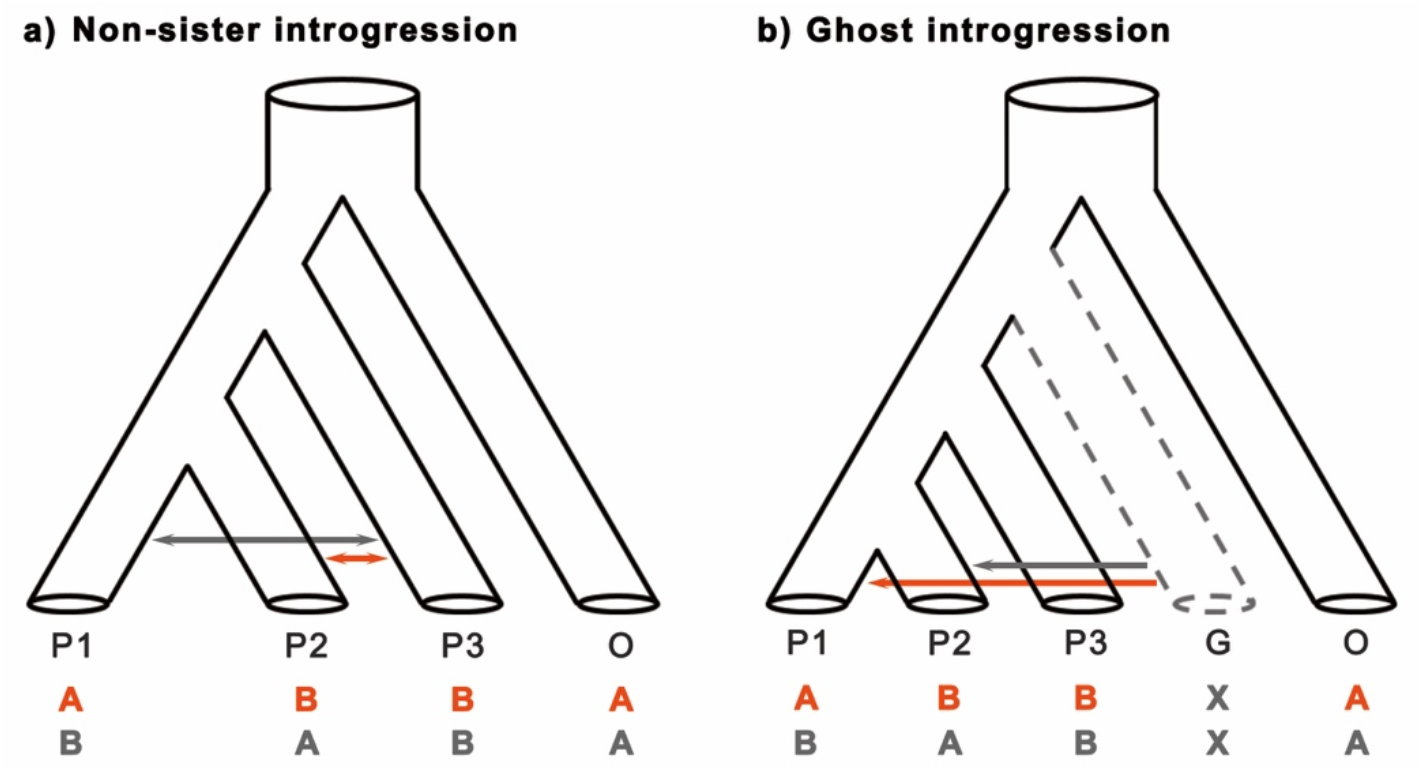
Introgression scenarios based on a ladder-like species tree of four extant lineages (ingroups: P1, P2, and P3; outgroup: O) and one ghost lineage (G). The non-sister introgression (a) and ghost introgression (b) events can result in a *D*-statistic significantly different from zero with a substantial excess of either ABBA or BABA patterns. The red or grey solid arrows indicate the introgression events between non-sister lineages or from the ghost lineage to one of the sister lineages.

Of note, the site patterns used in the *D*-statistic or alike do not provide enough information on their own to distinguish introgression from a nonsister lineage or a ghost lineage (Pang and Zhang 2023a). Therefore, extra analysis is clearly needed to distinguish the two kinds of introgression events (ingroup introgression between nonsister extant lineages or ghost introgression from an unsampled outgroup lineage) responsible for a significant *D*- statistic (Fig. 1). The full-likelihood program BPP (Bayesian Phylogenetics and Phylogeography), which uses a multispecies-coalescent-with-introgression (MSci) model (Flouri et al. 2020), is highly recommended for this purpose (Pang and Zhang 2023a). This program enables accurate estimation of the direction, timing, and magnitude of introgression under specified introgression models, and can compute marginal likelihoods to compare models across genomic datasets with single or multiple sequences from each species.

Therefore, by combining *D*-statistic and BPP analysis, researchers may be able to effectively and accurately identify introgression scenarios, particularly those involving ghost lineages.

Like most methods for inferring introgression, *D*-statistic also requires that the rooted species tree is known or can be inferred accurately; otherwise, it is likely to return meaningless results (Dagilis et al. 2022; Hibbins and Hahn 2022a; b). However, if extensive introgression occurs, it may be extremely hard, if not impossible, to retrieve the true bifurcation history of the species because phylogenetic inference based on DNA sequence alignments will likely produce species trees that reflect the reticulation history rather than the bifurcation history (Mallet et al. 2016; Jiao et al. 2020; Pang and Zhang 2023b). Dagilis et al. (2022) even warned against using the *D*-statistic or similar statistics altogether when the species tree is highly uncertain. In this context, Aguillon et al. (2022) have suggested that genomic regions with high gene density or low recombination rates can be used to reconstruct true phylogenetic relationships because these regions are unlikely to introgress due to potentially strong interference with gene function or linkage to deleterious alleles. However, genomic regions of low recombination or high gene density are not immune from the impact of introgression, not to mention that they can produce misleading tree topologies as a result of selection (Nater et al. 2015). Luckily, the phylogenetic signal from genome structure (gene content and order) appears reliable and relatively robust to introgression (Zhao et al. 2021; Ding et al. 2023). Allelic replacements due to cross-species gene flow can have a large influence on alignment-based phylogenetic inference, but will not impact the gene content and order of the recipient genome. Structural variants do have the potential to alter the gene content and order of the introgressed genome, but they are less likely to cross species boundary due to strong purifying selection against them (Zhou et al. 2019; Hämälä et al. 2021; Zhang et al. 2023). Thus, for taxa that have high-quality genome assemblies available, we may use microsynteny-based (Zhao et al. 2021) or gene-content-based (Delsuc et al. 2005; Pett et al. 2019) phylogenetic methods to accurately infer species trees in the presence of introgression (Ding et al. 2023).

The beaked hickory *Carya sinensis* Dode, also known as *Annamocarya sinensis* (Dode) Leroy, is a relic species of the walnut family (Juglandaceae), and its nut is generally characterized by a prominent apex, just like a bird’s beak (Supplementary Fig. S1). Some paleobotanists have posited that the fruits of *C. sinensis* display characteristics indicating an ancestral connection to both *Juglans* and *Carya*, leading to its classification as a living fossil of *Caryojuglans* (Merrill 1948) or *Juglandicarya* (Hu 1952). As a consequence, the phylogenetic placement of *C. sinensis* remains uncertain (Grauke et al. 1991): some researchers have considered it a monotypic genus due to its distinct morphological features (Leroy 1950; Kuang and Lu 1979; Lu et al. 1999), while others have included it within the genus *Carya* based on the early phylogenetic reconstructions using limited molecular markers (Manos and Stone 2001; Manos et al. 2007; Zhang et al. 2013; Xi et al. 2022). Currently, the beaked hickory grows in dense, lowland forests located in southwestern China and northern Vietnam where there is very little light penetrating the forest floor. The genus *Carya* consists of 17-19 species, with disjointed distributions in eastern Asia (EA) and North America (NA) (Grauke et al. 2016; Kozlowski et al. 2018; Zhang et al. 2022a). Due to the uncertainty surrounding the classification and systematic position of the beaked hickory in relation to *Carya*, we suspect that ancient introgression might have occurred from a distant unknown relative, resulting in the ancestral characteristics of the beaked hickory.

Here, we focus on the possibility of introgression between *C. sinensis* and other diploid *Carya* species, including ghost lineages derived from the common ancestor of *C. sinensis* and *Carya*. First, we generated two new high-quality chromosome-level genomes for *C. sinensis* (beaked hickory) and *C. cathayensis* (Chinese hickory), and downloaded the previously published genomes of *C. illinoinensis* (pecan) (Lovell et al. 2021) and *Pterocarya stenoptera* (Chinese wingnut) (Zhang et al. 2022b), which enable us to obtain a reliable species tree topology [((P1: *C. sinensis*, P2: *C. cathayensis*), P3: *C. illinoinensis*), O: *P. stenoptera*] with whole-genome microsynteny (Zhao et al. 2021) and gene content (Delsuc et al. 2005; Pett et al. 2019) methods. Second, we combine the *D*-statistic with a full-likelihood method for detecting gene flow — the MSci model in BPP (Flouri et al. 2020) — to determine whether the beaked hickory experiences ancient introgression from an unsampled ghost taxon. Third, we use 7,398 single-copy nuclear loci to reconstruct the phylogenetic network of *C. sinensis* and 11 diploid *Carya* species (6 EA *Carya* species and 5 NA *Carya* species) with PhyloNet (Yu and Nakhleh 2015; Wen et al. 2018), finding that about a quarter of the *C. sinensis* nuclear genome is derived from an unknown ancestral lineage of *Carya*. This is further reinforced by the results obtained from the BPP analysis of a subset of 773 loci from the complete dataset, due to heavy computational costs. Finally, recognizing that VolcanoFinder (Setter et al. 2020) may provide independent evidence for ghost introgression because this program is capable of detecting signatures of adaptive introgression from unknown donors, we applied this program to analyze the whole-genome resequencing data from 43 individuals of *C. sinensis*. Our objective was not only to provide additional evidence supporting the presence of ghost introgression in the beaked hickory but also to identify potential candidate introgressed genes that contribute to adaptation to the sustained shading conditions commonly found in dense forests.

## MATERIALS AND METHODS

Fresh young leaves were sampled from one wild plant of *C. sinensis* (Malipo County, Yunnan Province, China, 23°7′49.38″N, 104°51′30.59″E) and of *C. cathayensis* (Mt. Tianmushan, Zhejiang Province, China, 30°19′31.62″N, 119°26′27.79″E) and PacBio single-molecule real-time long reads and chromosome conformation capture (Hi-C) technique were used to assemble the chromosome-scale genomes for the two species. The 43 individuals of *C. sinensis* and one individual from each of the 11 diploid *Carya* species were also collected for whole genome resequencing on the Illumina NovaSeq 6000 platform.

We investigate the phylogenetic relationships between *C. sinensis* and the genus *Carya*, and conduct the introgression tests and subsequent analyses based on the *de novo* genome sequences and whole-genome resequencing dataset. First, we carry out a comparative genomic analysis of *C. sinensis*, *C. cathayensis* (from EA *Carya*), *C. illinoinensis* (from NA *Carya*) (Lovell et al. 2021) using MCScanX (Wang et al. 2012) and KaKs_Calculator 2.0 (Wang et al. 2010). Second, we infer the whole-genome tree of *C. sinensis*, *C. cathayensis* and *C. illinoinensis*, with *P. stenoptera* (Zhang et al. 2022b) serving as the outgroup, using three methods: ASTRAL-Pro (Zhang et al. 2020), whole-genome microsynteny (Zhao et al. 2021), and gene content (Delsuc et al. 2005; Pett et al. 2019). In addition, we employ ASTRAL v5.7.4 (Mirarab et al. 2014) to construct a species tree for *C. sinensis*, six EA *Carya* species, and five NA *Carya* species from their dataset of whole-genome resequencing. Third, the *D*-statistic is applied to the dataset consisting of 11,269 single-copy genes from one individual per species, for the purpose of conducting an introgression test based on 30 possible rooted quartets of [((P1: *C. sinensis*, P2: EA *Carya* species), P3: NA *Carya* species), O: *P. stenoptera*]. Then, the MSci model implemented in the BPP v4.6.2 program (Flouri et al. 2020) is utilized with three triplets of hickory species (excluding the outgroup *P. stenoptera*) using 2,543 single-copy genes to ascertain the introgression scenario (Fig. 1) that is truly responsible for the significant *D*-statistic observed in all of the aforementioned 30 species quartets (see Results). These 2,543 loci are carefully selected from the pool of the previously mentioned pool of 11,269 single-copy genes, satisfying the following filtering criteria: (i) absence of recombination events, (ii) less than 20% gaps and less than 50% missing data in each sequence, and (iii) sequence alignment lengths ranging between 500 and 1,000 base pairs (bps). Fourth, the PhyloNet/MPL (Yu and Nakhleh 2015; Wen et al. 2018) is employed to reconstruct phylogenetic networks for 12 taxa, which include *C. sinensis*, six EA *Carya* species, and five NA *Carya* species, from 7,398 gene trees under a maximum pseudo-likelihood estimation method. Each of the 7,398 single-copy genes used in network reconstructions meets the filtering constraints (i) and (ii) and sequence lengths ranging from 300 to 2,000 bps. Additionally, to estimate the timing of ghost introgression in *C. sinensis*, the MSci model and 773 single-copy nuclear loci were employed using BPP v4.6.2 (Flouri et al. 2020). The reduced 773 loci were selected from the collection of 7,398 single-copy genes, with a specific focus on sequences ranging from 800 to 1,000 bps in length, in light of the computational limitations related to BPP analysis. Finally, we employ VolcanoFinder v1.0 (Setter et al. 2020) to analyze the resequencing data from 43 individuals of *C. sinensis*, aiming to identify instances of adaptive introgression in this species. Detailed information on materials and methods with associated references are available in the Supplementary Materials.

## RESULTS

### Genome Features and Comparative Genomic Analysis

We integrated three different sequencing technologies (Illumina NovaSeq 6000 platform (Illumina), single-molecule real-time long reads (PacBio), and chromosome conformation capture (Hi-C)) to *de novo* assemble the whole genomes of *C. sinensis* and *C. cathayensis* (Supplementary Text, Fig. 2a and Supplementary Fig. S2). In total, 235.51 Gb (∼371×) and 207.53 Gb (∼280×) of raw data from two species were sequenced, with the estimated genome size of 634.52 Mb and 742.32 Mb for *C. sinensis* and *C. cathayensis*, respectively. The resulting whole genomes were assembled at the chromosomal level, yielding a total length of 623.16 Mb (scaffold N50 of 38.85 Mb) for *C. sinensis* and 698.09 Mb (scaffold N50 of 43.49 Mb) for *C. cathayensis* (Supplementary Table S1). A total of 94.5% and 93.9% of all universal single-copy orthologs in BUSCO’s embryophyta odb9 database were present and complete in *C. sinensis* and *C. cathayensis*, respectively (Supplementary Table S2). These results suggested that the two genomes of *C. sinensis* and *C. cathayensis* possess a high level of continuity, completeness, and accuracy. Additionally, we predicted 35,370 protein-coding genes in the *C. sinensis* genome and 36,722 in the *C. cathayensis* genome (Supplementary Table S1). More than 90.33% of genes in *C. sinensis* and 88.76% in *C. cathayensis* could be assigned to entries in four functional databases of GO, KEGG, NR, and UniProt-TrEMBL (Supplementary Table S3). Through syntenic analysis, we observed a substantial collinearity between the genomes of *C. sinensis, C. cathayensis*, and *C. illinoinensis* (Fig. 2b). The distribution of *K*_s_ values among paralogous gene pairs within the three *Carya* species exhibited two distinct peaks at 0.31 and 1.45, suggesting two whole-genome duplication (WGD) events (Fig. 2c). Furthermore, the *K*_s_ distribution of orthologous gene pairs in three different species pairs showed striking similarities, with the *C. sinensis* - *C. cathayensis* pair displaying the lowest peak value (Supplementary Fig. S3).

**FIGURE 2.**
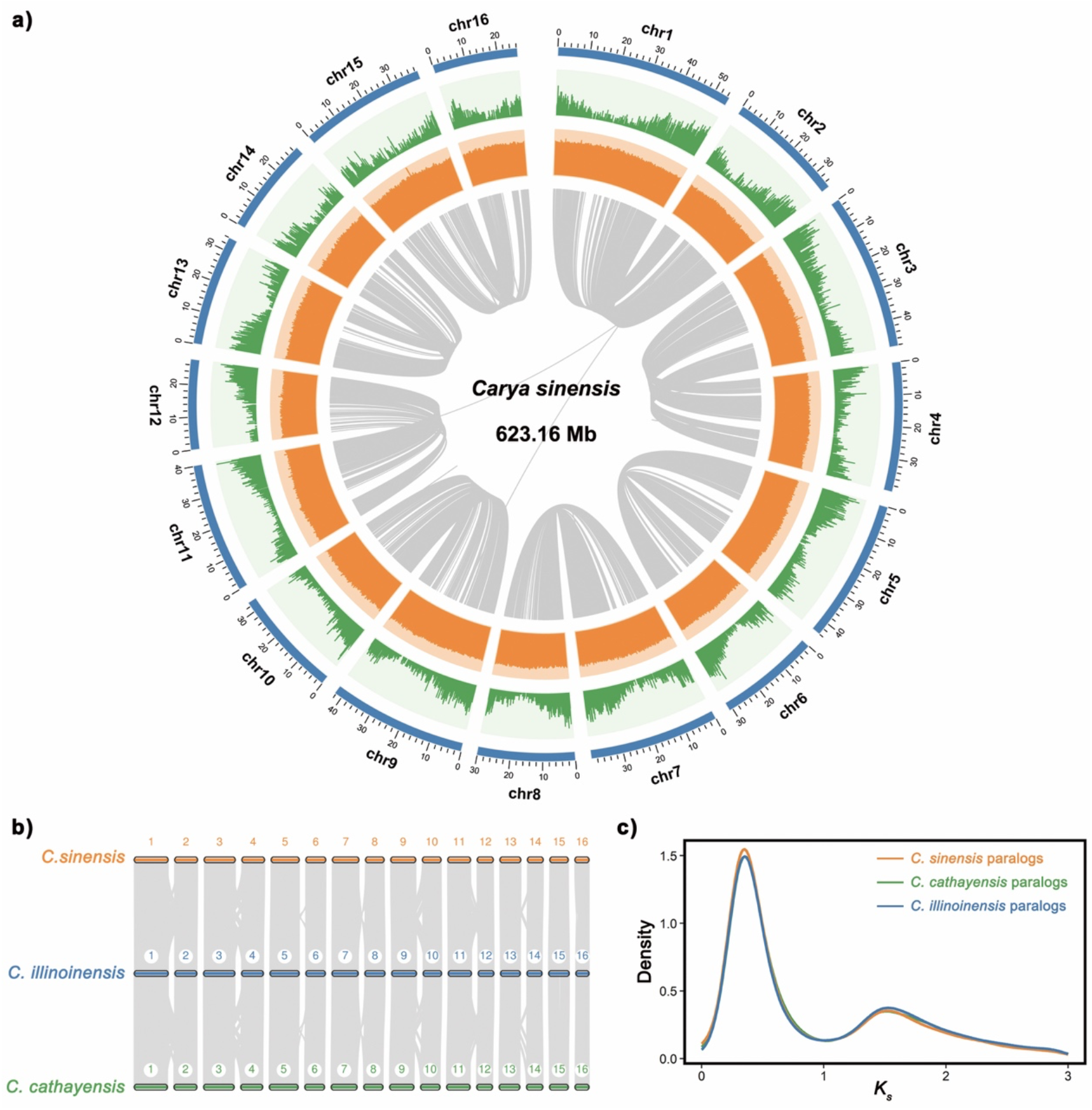
Genome features of *Carya sinensis* and comparative genomic analysis with other two hickory species. **a)** Overview of the *C. sinensis* genome. Different tracks (moving inward) denote (i) 16 chromosomes; (ii) gene density in 100 kb stepping windows (minimum-maximum, 0–24); (iii) GC content in 100 bp stepping windows (minimum-maximum, 0.2–0.5); (iv) identified syntenic blocks. **b)** MCScanX identified synteny blocks between *C. sinensis*, *C. cathayensis,* and *C. illinoinensis*. **c)** The *K*_s_ (synonymous substitution rate) distributions for syntenic gene pairs within *C. sinensis*, *C. cathayensis,* and *C. illinoinensis*.

### Species Tree Reconstruction and Introgression Test

Three methods of species tree construction, i.e. ASTRAL-Pro (Zhang et al. 2020) based on DNA-sequence alignment, as well as whole-genome microsynteny (Zhao et al. 2021) and gene content (Pett et al. 2019) methods based on genome structure, consistently yielded the topology [((*C. sinensis*, *C. cathayensis*), *C. illinoinensis*), *P. stenoptera*] (Supplementary Text and Supplementary Fig. S4). Additionally, we utilized 7,398 gene trees of single-copy genes obtained from one re-sequenced individual of *C. sinensis* and 11 diploid *Carya* species, with *P. stenoptera* as the outgroup (Supplementary Text and Supplementary Tables S4-S6), to reconstruct the phylogenetic tree of hickory species using ASTRAL v5.7.4 (Mirarab et al. 2014). The resulting tree revealed that *C. sinensis* and six EA *Carya* species formed a clade, which is sister to a clade consisting of five NA *Carya* species. This finding aligns with the whole-genome tree of the three species constructed using ASTRAL-Pro, gene content, and microsynteny methods, and corresponds to the biogeographic disjunction between eastern Asia (EA) and North America (NA) observed in extant *Carya* species (Supplementary Fig. S5).

The introgression tests were performed by applying the *D*-statistic method to 30 rooted quartets of [((P1: *C. sinensis*, P2: EA *Carya* species), P3: NA *Carya* species), O: *P. stenoptera*] (Supplementary Text). The *D*-statistic results for all 30 quartets exhibited significant positive values (Supplementary Table S7), indicating the occurrence of introgression events. In addition to the previously mentioned non-sister introgression (Fig. 1a) and ghost introgression from an outgroup lineage (Fig. 1b), which are likely to yield positive *D* values, the possibility of ghost introgression originating from the ingroup lineage should also be considered. Consequently, we established six introgression models, potentially involving EA *Carya* species (P2) and NA *Carya* species (P3) (Supplementary Fig. S6a-c), or encompassing the introgression from an ingroup ghost lineage into NA *Carya* species (Supplementary Fig. S6d), or involving the introgression from an ingroup ghost lineage into EA *Carya* species (Supplementary Fig. S6e), or incorporating the introgression from an outgroup ghost lineage into *C. sinensis* (Supplementary Fig. S6f). Then, the BPP v4.6.2 (Flouri et al. 2020) was used to compare the six candidate introgression scenarios (Supplementary Fig. S6) for three representative triplets, employing a total of 2,543 single-copy genes owing to the considerable computational loads involved (Supplementary Text). For the triplet ((*C. sinensis*, *C. cathayensis*), *C. illinoinensis*), the log marginal likelihood was -3,138,311 for the outgroup ghost introgression scenario (model 6 in Supplementary Fig. S6f), which was higher than the marginal likelihoods of other five introgression scenarios (models 1-5 in Supplementary Fig. S6a-e). The BPP analysis of two additional triplets, ((*C. sinensis*, *C. tonkinensis*), *C. illinoinensis*) and ((*C. sinensis*, *C. kweichowensis*), *C. cordiformis*), also strongly supported introgression from outgroup ghost lineage into *C. sinensis* (Supplementary Table S8). Therefore, the combined results of the *D*-statistic and BPP analyses provide strong evidence supporting the occurrence of introgression from an ancestral ghost lineage into *C. sinensis*.

### Phylogenetic Network and Divergence Time Inference Based on Whole-Genome Resequencing Data

The PhyloNet analyses (Wen et al. 2018), which incorporate incomplete lineage sorting and hybridization, classified the species into two major clades: EA *Carya* and NA *Carya* (Supplementary Text, Supplementary Figs. S7 and S8), consistent with the ASTRAL results without considering gene flow (Supplementary Figs. S4 and S5). In all instances allowing two or three hybridization events, *C. sinensis* was invariably identified as a reticulate node (Supplementary Figs. S7 and S8), with a genomic contribution of 29% (in the case of two reticulations) or 26% (in three reticulations) from the ancestral ghost lineage. These findings are substantiated by the aforementioned BPP analysis, which examined three triplets and employed a dataset consisting of 2,543 single-copy genes (Supplementary Table S8). In addition, the phylogenetic network analysis suggests that *C. hunanensis* has a hybrid origin (Supplementary Fig. S7), further exploration of this intriguing finding will be undertaken in future studies.

To estimate the divergence time, we applied the BPP implementation of the MSci model (Flouri et al. 2020) to the reduced genomic dataset of 773 single-copy nuclear loci for 12 taxa (Supplementary Text). Our dating estimates indicate that the ghost lineage diverged from the common ancestor of hickories, including *C. sinensis* and the other 11 diploid *Carya* species, approximately 5.42 million years ago (95% highest posterior density [HPD] interval: 4.97- 5.89 Ma) (Fig. 3). Furthermore, we estimated that the introgression from the ghost lineage into *C. sinensis* occurred around 2.72 Ma (95% HPD interval: 2.24-3.03 Ma), with an inheritance probability of approximately 0.22 (95% HPD interval: 0.15-0.28) (Fig. 3). Our results also revealed that the split between the extant EA (including *C. sinensis*) and NA *Carya* clades happened around 3.96 Ma (95% HPD interval: 3.87-4.05 Ma), with subsequent diversification occurring at 2.96 Ma (95% HPD interval: 2.86-3.06 Ma) for EA *Carya* and 2.01 Ma (95% HPD interval: 1.95-2.07 Ma) for NA *Carya* (Fig. 3). Disregarding introgression, similar divergence time estimates were obtained from the BPP analysis using the multispecies coalescent (MSC) model (Supplementary Fig. S9).

**FIGURE 3.**
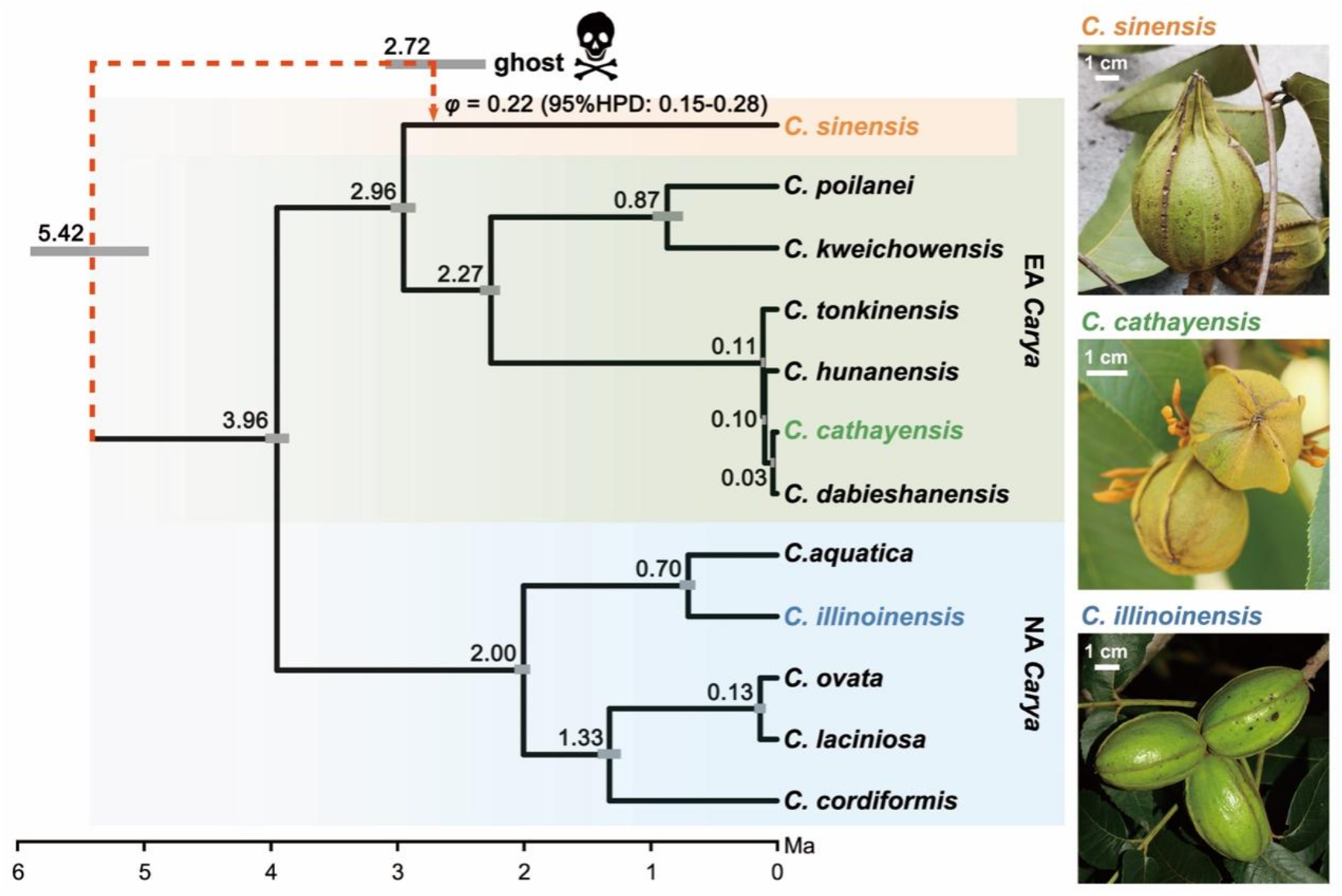
The divergence time estimates of the ghost introgression scenario from the *Carya sinensis* and the other 11 diploid *Carya* species under the MSci model using the software BPP and the reduced dataset of 773 loci. Node heights are posterior means under the MSci model with the absolute divergence times. The red dotted arrow is shown by ghost lineage along with their corresponding introgression probabilities from the MSci model. Posterior means and 95% HPD CIs for the ghost introgression donor and recipient nodes are shown above their vertices to improve visibility. The three fruit morphology images along the right show the *Carya sinensis*, *C. cathayensis* (represents EA *Carya* species), and *C. illinoinensis* (represents NA *Carya* species), respectively.

### Possibly Ghost Introgressed Genes and Adaptive Signals

To examine whether ghost introgression in *C. sinensis* did occur, we conducted comprehensive genome-wide scans of adaptive introgression using VolcanoFinder v1.0 (Setter et al. 2020) based on polymorphism data from 43 individuals of *C. sinensis* (Supplementary Text). Our analysis focused on identifying 44 20-kb blocks exhibiting composite-likelihood ratio (CLR) values exceeding 100, within which we identified a total of 36 candidate genes that could potentially be the result of ghost introgression (Supplementary Tables S9 and S10). Furthermore, by performing Gene Ontology (GO) enrichment analysis on these genes, we revealed their involvement in various biological processes such as the positive regulation of circadian rhythm, red light and far-red light phototransduction, the far-red light signaling pathway, and defense response (Supplementary Fig. S10, Supplementary Tables S9 and S10). Notably, the detected genes included FHY3 and FAR1 transcription factors, which play crucial roles in phytochrome A-mediated light signaling and far-red light signaling in *Arabidopsis* (Stirnberg et al. 2012; Xie et al. 2020). These genes are recognized for their ability to trigger reduced branching and accelerated shoot elongation as a response to continuous shading, potentially playing a role in the adaptation of *C. sinensis* to thrive under plant canopy shade in subtropical and tropical forests.

## DISCUSSION

### Phylogenetic Position of Carya sinensis Within the Genus Carya

A robust phylogenetic framework is crucial for comprehending the evolutionary history and species interactions. The construction of species trees for the three genomes, using three different methods, namely ASTRAL-Pro (Zhang et al. 2020), whole-genome microsynteny (Zhao et al. 2021), and gene content (Delsuc et al. 2005; Pett et al. 2019), consistently revealed a sister relationship between *C. sinensis* and *C. cathayensis* (Supplementary Figs. S3 and S4). Additionally, the species tree derived from re-sequenced genomes of *C. sinensis*, six EA *Carya* species, and five NA *Carya* species showed that *C. sinensis* is the sister taxon to the EA *Carya* Clade (Supplementary Fig. S5). Hence, it is suggested that *C. sinensis* should be classified as a member of the *Carya* genus rather than as a monotypic genus called *Annamocarya*. Despite exhibiting characteristics such as semi-evergreen plants, entire leaflets, and large fruit with keeled segments of the thick husk (Supplementary Fig. S1) (Leroy 1950; Manning 1963; Lu et al. 1999), the fundamental features of *C. sinensis* align more closely with the *Carya* genus.

### Genomic Evidence of Ghost Introgression in Carya sinensis

Three lines of evidence support the scenario of ghost introgression, with *C. sinensis* acting as the recipient of gene flow. First, the presence of introgression detected by the *D*- statistic with 30 rooted quartets of [((P1: *C. sinensis*, P2: EA *Carya* species), P3: NA *Carya* species), O: *P. stenoptera*] (Supplementary Table S7) was confirmed to be from an outgroup ghost lineage through comparison of six introgression models (Supplementary Fig. S6) using the full-likelihood BPP program. Second, the network estimated by PhyloNet/MPL (Yu and Nakhleh 2015; Wen et al. 2018) provides additional support for the presence of ghost introgression in *C. sinensis* (Supplementary Figs. S7 and S8). The estimated inheritance probabilities (*γ*) of 0.29 or 0.26 in the networks allowing two or three reticulation events respectively (Supplementary Figs. S7 and S8) demonstrate a resemblance to the estimate obtained using the BPP program with the MSci model (Flouri et al. 2020) incorporating *C. sinensis* and 11 other diploid *Carya* species (Fig. 3). Third, the analysis conducted by VolcanoFinder (Setter et al. 2020) identified 36 possible genes involved in ghost introgression, including FHY3 and FAR1 transcription factors (Supplementary Tables S9 and

S10). These genes hold the potential to shed light on the adaptive mechanisms employed by *C. sinensis* in adapting to the dim light conditions prevalent in subtropical and tropical forest canopies (Supplementary Fig. S10).

### Advantages of the Combined Method of D-statistic and BPP

Our analysis emphasizes the advantages of combining the *D*-statistic and BPP methods, which collectively facilitate the precise identification of introgression types. The methods are relatively simple to compute and interpret. The *D*-statistic only requires a single sequence from each of the four taxa, and it can robustly detect the presence of introgression across various scenarios (Martin et al. 2015; Zheng and Janke 2018; Hahn and Hibbins 2019; Hibbins and Hahn 2022a). Because the full-likelihood BPP method takes into account the information of topologies and branch lengths in gene trees, it can discriminate between ghost introgression and introgressions between non-sister species that are indistinguishable by the *D*-statistic method (Pang and Zhang 2023a). Furthermore, BPP can accurately estimate the timing of introgression and species divergence times, even when a single diploid individual (or two haploid sequences) per species is sampled (Tiley et al. 2023). Therefore, when the *D*- statistic indicates the presence of introgression, BPP can be employed to confirm the more likely introgression scenario. However, it is important to note certain caveats. BPP assumes a JC69 mutation model (Jukes and Cantor 1969), which may not be appropriate for distantly related species (Yang 2015; Flouri et al. 2018; 2020). Moreover, the loci used in the BPP program should be independent and free of recombination within a locus (Yang 2015; Flouri et al. 2018).

Although our findings in PhyloNet are consistent with the recommendation of Solis-Lemus and Ane (2016) that the introgression edge should be assigned to the edge with a minor contribution, previous studies have demonstrated that introgression can have a significant impact on the majority of the genome (Fontaine et al. 2015; Li et al. 2019; Forsythe et al. 2020; Hibbins and Hahn 2022b). Therefore, interpreting the phylogenetic networks should be cautious with regard to speciation and introgression edge. Compared to PhyloNet/MPL, which relies on the probabilistic inference of phylogenetic networks based on gene tree topologies, the full-likelihood method BPP capitalizes on both gene tree topologies and branch lengths, enabling a more accurate determination of specific scenarios of introgression.

### Implications for the Biogeography of Hickories

In our BPP analysis of ghost introgression in the beaked hickory, the age of the donor lineage was estimated to be 5.42 Ma, indicating that the extinct taxon had a presence in eastern Asia (EA) during this period, presenting an opportunity for genetic material to introgress into *C. sinensis* at 2.72 Ma (Fig. 3). According to Manchester (1987) and Zhang et al. (2013), the genus *Carya* probably originated in North America (NA) during the early Paleocene, and subsequently migrated to Europe via the North Atlantic land bridge (NALB) during the Paleocene and Eocene. Then, the genus *Carya* may have entered Asia either from Europe as the Turgai seaway receded or via the Bering land bridge (BLB) during the late Miocene and Pliocene. However, our dating analysis using BPP estimated that the crown age of extant *Carya* species from East Asia (EA) and North America (NA) was approximately 3.96 Ma. Therefore, we propose the additional possibility that extant *Carya* species in North America (NA) migrated back from Asia via the BLB (Supplementary Fig. S11), while the original hickory species in North America (NA) and Europe went extinct due to the cooling period across the high latitude areas of the Northern Hemisphere, beginning in the Miocene (Wolfe 1975). The back-migration likely occurred around 3.96 Ma as estimated by our BPP analysis. Evidence from the morphological change of bud scales in *Carya* may support our back-migration hypothesis, as EA *Carya* species have naked buds and atypical bud scales, which are considered primitive, while NA *Carya* species have imbricated and valvated bud scales, which are considered an evolutionary adaptation to temperate climate (Chang and Lu 1979).

## CONCLUSION

As advised in several pertinent studies, it is crucial to consider the potential influence of ghost lineages when discerning the distinct types of introgression (Ottenburghs 2020; Hibbins and Hahn 2022a; Tricou et al. 2022a; b). In this study, we elucidated the systematic position of *C. sinensis* within the genus *Carya* by reporting two newly assembled genomes of *C. sinensis* and *C. cathayensis* and resequencing data from 54 individuals of 12 hickory species. Our combined use of the *D*-statistic and BPP methods provides strong evidence supporting the existence of ghost introgression in *C. sinensis*, while simultaneously refining the understanding of *Carya*’s biogeography via divergence estimates. Overall, our findings endorse the notion that genetic imprints of introgression may indeed originate from extinct or unsampled “ghost” lineages, thereby offering new insights into the evolutionary history of the relevant lineages.

## DATA AVAILABILITY

The raw sequence data reported in this paper have been deposited in the Genome Sequence Archive in National Genomics Data Center, China National Center for Bioinformation/Beijing Institute of Genomics, Chinese Academy of Sciences (GSA: CRA011421) that are publicly accessible at https://ngdc.cncb.ac.cn/gsa. The final genome assembly and genome annotation information has also been available on the website (http://cmb.bnu.edu.cn/juglans).

## ACKNOWLEDGMENTS

We are grateful to Patrick Thompson (Davis Arboretum, the United States), Joke Ossaer (Foundation Arboretum Wespelaar, Belgium), and Shuai Liao (South China Botanical Garden, Chinese Academy of Sciences) for assistance with sampling, and Chao-Yi Deng (Agriculture and Forestry Research Inst. of southwestern Guizhou Province) and Xin-Xin Zhu (Xinyang Normal University) for providing photos of *Carya sinensis* and *C. illinoinensis*, respectively.

