## Supplementary Materials for "Uncovering Ghost Introgression Through Genomic Analysis of a Distinct East Asian Hickory Species"

### Supplementary Text

#### *Sample Collection,* *Genome Sequencing and Assemblies*

Fresh young leaves of *Carya sinensis* (Malipo County, Yunnan Province, China, 23°7′49.38″N, 104°51′30.59″E) and *C. cathayensis* (Mt. Tianmushan, Zhejiang Province, China, 30°19′31.62″N, 119°26′27.79″E) were sampled from two wild plants and immediately stored in liquid nitrogen. All samples were sent to NovoGene (Beijing, China) for genomic sequencing. The genomic libraries with 20 kb insertions were constructed and sequenced for the single-molecule long reads using a PacBio-sequel platform. In addition, a Hi-C experiment was orderly performed as follows: cell cross-linking, endonuclease digestion, terminal repair, cyclization, DNA purification and capture. Then the libraries were constructed and sequenced using the Illumina NovaSeq 6000 platform. For the Illumina short reads, the DNA libraries with 350 bp insert sizes were constructed and sequenced using an Illumina NovaSeq 6000 platform. All sequencing raw reads were processed to remove adaptors, low-quality bases, and possible contaminated sequences.

To estimate the genome size of *C. sinensis* and *C. cathayensis*, the 150 bp paired-end reads were surveyed with 17 bp *k*-mer frequencies using Jellyfish v2.3 (Marcais and Kingsford 2011). The resulting *k*-mer count histograms were then evaluated using GenomeScope 2.0 (Ranallo-Benavidez et al. 2020).

After correcting reads by Canu v1.8 (Koren et al. 2017), the PacBio single-molecule long reads were preassembled in the following procedures. First, the initial contigs assembly was generated via Falcon v3.1 (Chin et al. 2016) after error correction. Second, contigs were polished by mapping the Illumina short reads through QUIVER (Chin et al. 2013) and were further corrected by Pilon v1.22 (Walker et al. 2014) with default parameters to obtain the contig-level assembly reference. The GC content and sequencing coverage analyses were applied to evaluate the presence of contamination. To assess the assembly quality, the embryophyta_odb9 database of 1,440 conserved plant genes was searched against the genome sequences through BUSCO v3.0.2 (Simao et al. 2015). The Hi-C sequencing clean data were mapped to contig sequences by BWA v0.7.17 (Li and Durbin 2009), and the uniquely mapped read pairs were retained and further filtered by HiC-Pro v2.11.1 (Servant et al. 2015). Only the valid interaction pairs were used to cluster, order, and orient the contigs onto pseudo-chromosomes by LACHESIS (Burton et al. 2013). Finally, the errors of placement and orientation exhibiting obvious discrete chromatin interaction patterns were manually adjusted.

#### *Gene Prediction and Functional Annotation*

RepeatMasker v4.0.7 (http://www.repeatmasker.org) and RepeatModeler v1.0.8 (Benson 1999) were used to annotate the repeat sequences. To annotate the genomes of *C. sinensis* and *C. cathayensis*, the combination of *ab initio* prediction, homology-based inference, and transcripts from RNA sequencing (RNA-Seq) was conducted (the texts below show the analysis process in *C. sinensis*, and the same method was also used in *C. cathayensis*). The RNA-seq data sequenced from the leaf and flower bud tissues were mapped to the *C. sinensis* genome using Hisat2 v2.0.4 (Kim et al. 2015). Alignments were further assembled into the *de novo* and genome-guided transcripts by StringTie v1.3.0 (Pertea et al. 2016). All assembled transcript sequences were run by PASA v2.1.0 (Haas et al. 2003) for gene structure annotation. The AUGUSTUS v3.2.3 (Stanke et al. 2006) was applied to produce *ab initio* evidence. The proteins of *Arabidopsis thaliana*, *Betula pendula*, *Castanea mollissima*, *Cucumis sativus*, and *Juglans regia* from the NCBI database, and proteins of all plants from the Swiss-Prot database, were collected for homologous prediction mainly by GeneWise v2.4.1 (Birney et al. 2004). All gene models were integrated using EVM v1.1.1 (Haas et al. 2008), and then filtered according to the following four criteria: no start codon or stop codon, length of CDS being not a multiplier of 3 or less than 150 bp, containing premature-stop codon, and EVM score lower than 4000. The final gene sets were functionally annotated using BLASTP (minimum mapping length of 50 bp, minimum identity of 50%, minimum coverage of 50%, and minimum e-value of 1e-5) against the NCBI NR, UniProt-TrEMBL, and KEGG databases. Gene Ontology (GO) annotation was performed using Blast2GO (Conesa et al. 2005), and pathway annotation was performed using KAAS (Moriya et al. 2007).

#### *Whole-Genome Duplication and Synteny Analysis*

To identify the whole-genome duplication (WGD) event, the collinear blocks within *C. sinensis*, *C. cathayensis,* and *C. illinoinensis* (Lovell et al. 2021) genomes were identified by MCScanX (Wang et al. 2012) using default parameters, respectively. The intraspecific collinear blocks were visualized by TBtools (Chen et al. 2020). For collinear genes (anchor genes) of each collinear block, the synonymous substitution rate (*Ks*) was estimated using the Gamma-MYN algorithm (Wang et al. 2009) in KaKs_Calculator 2.0 (Wang et al. 2010). The distribution of *K_s_* was plotted using ggplot2 in R. The collinear blocks of any two species among *C. sinensis*, *C. cathayensis*, and *C. illinoinensis* were identified using the MCScanX (Wang et al. 2012) implemented in Python (https://github.com/tanghaibao/jcvi/wiki/MCscan-(Python-version)).

#### *Genome Re-sequencing, Read Mapping, and Variant Calling*

To perform the phylogenetic network and adaptive introgression analysis, 43 individuals of *C. sinensis* from most of the known locations across its distribution range and 11 individuals representing 11 diploid *Carya* species were collected (Supplementary Tables S4-S6)). For each individual, 4-6 flesh leaflets were dried in silica gel and stored at room temperature. DNA was extracted according to the manufacturer’s protocol using the HP Plant DNA Kit D2485-02 (Omega Bio-Tek). Whole-genome re-sequencing was performed on Illumina NovaSeq 6000 instruments by NovoGene (Beijing, China). All individuals were expected to sequence to a depth of 30× with paired-end libraries of 350 bp insert size and read length of 150 bp. Voucher specimens for these species were deposited in the Herbarium of the College of Life Science, Beijing Normal University (BNU).

The raw reads were quality controlled and trimmed for adapters using FastQC and Trimmomatic v0.32 (Bolger et al. 2014), respectively. Then the high-quality clean reads of each species were mapped to its closely related reference genome using the BWA-MEM algorithm of BWA v0.7.12 with default settings (Li 2013). In brief, 43 individuals of *C. sinensis*, six individuals of six species of eastern Asian *Carya* (EA *Carya*)*,* and five individuals of five species of North American *Carya* (NA *Carya*) were mapped onto the *C. sinensis*, *C. cathayensis,* and *C. illinoinensis* genome, respectively. SAMtools v0.1.19 (Li 2011) was used to retain only uniquely mapped and properly paired reads and to convert the Sequence Alignment Map (SAM) to a Binary Alignment Map (BAM) format file. The SENTIEON DNAseq software package v201808.08 (Weber et al. 2016) was subsequently used to remove duplicate reads, realign indels, and call single nucleotide polymorphisms (SNPs). The high quality of genome-wide SNPs was controlled by the following criteria: (i) sites with depth less than one-third or more than two-fold average depth per individual, non-bi-allelic sites, and sites with missing data were removed; (ii) the heterozygous genotypes were called if the proportion of the non-reference allele was falling inside the 20–80% when the sequencing depth is >20× (Nielsen et al. 2011), or if the proportion of the non-reference allele was falling inside the 10–90% when the sequencing depth is ≤20× but >10× (Zhang et al. 2019); otherwise, a homozygous genotype would be called. After serious specific filtering steps, a total of 24,751,236 biallelic SNPs for 43 individuals of species *C. sinensis* were obtained. For the other 11 *Carya* species, the high-confidence SNPs and invariant sites were used to reconstruct consensus genomes.

#### *Construct Phylogenetic Trees Based on DNA Sequence Alignments or Genome-Structural Data*

The accurate species tree is the primary prerequisite for detecting introgression. Some methods for species tree inference such as those based on minimum gene tree node heights (Fontaine et al. 2015; Forsythe et al. 2020) have been proven to be unreliable and depend on introgression rates across the genome (Hibbins and Hahn 2022). Therefore, we apply genome structural data from whole-genome microsynteny (Zhao et al. 2021) and gene content (Pett et al. 2019), to reconstruct the phylogeny of *C. sinensis*, *C. cathayensis*, and *C. illinoinensis* (Lovell et al. 2021), with *Pterocarya stenoptera* (Zhang et al. 2022) serving as the outgroup. As a reference, we also use the conventional DNA-sequence-based phylogenetic method of ASTRAL-Pro (Zhang et al. 2020) to obtain a species tree for the three hickory species and the outgroup *P. stenoptera*.

For the DNA-sequence-alignment method, we identified the gene families using OrthoFinder v2.4.0 (Emms and Kelly 2019) based on protein-coding sequences from four species. To avoid multiple sequence alignment errors introduced by too large gene families (Thompson et al. 2011), we retained only those gene families with a total number of genes no more than 24, as these species experienced *γ*-WGT (whole-genome triplication) and 'juglandoid WGD' (Ding et al. 2023). The protein sequences of each gene family were aligned by MAFFT v7.475 (Katoh and Standley 2013) and then converted into corresponding codon alignments by PAL2NAL v14 (Suyama et al. 2006). IQ-TREE v2.1.2 (Minh et al. 2020) was employed to construct the maximum likelihood (ML) tree for each gene family with the parameters setting “-m MFP -B 1000”. Then the ASTRAL-Pro v1.1.6 (Zhang et al. 2020) was used to infer the species tree based on gene family trees under the MSC model and birth-death (GDL) model (Arvestad et al. 2009).

For whole-genome microsynteny-based phylogenetic reconstruction, we used synteny matrix representation with a likelihood (Syn-MRL) pipeline (Zhao et al. 2021). Following Zhao et al. (2021), BLASTP (Camacho et al. 2009) was used to perform the searches for all potential inter- and intra-species homologous gene pairs (default parameters). MCScanX (Wang et al. 2012) was used to detect the synteny blocks of inter- and intra-species with the recommended parameters of A_5_G_25_ (A: the minimum number of anchor pairs required to call a collinear block, G: maximum number of intervening genes between two (adjacent) anchor pairs in collinear blocks). In the synteny network, nodes are collinear genes in collinear blocks, and edges are used to connect collinear gene pairs. Using the Infomap algorithm v1.6.0 (https://github.com/mapequation/infomap), we detected synteny clusters within the map equation framework, setting the two-level partitioning mode with ten trials (--clu -N 10 -2). The entire network was assigned to 17,107 clusters using the Infomap algorithm v1.6.0. These clusters were transformed into a binary presence-absence data matrix where rows and columns represent species and clusters, respectively. The resulting matrix of 4 rows × 17,107 columns was used to infer a phylogeny with IQ-TREE v2.1.2 (Minh et al. 2020) under the Mk + R + FO model.

For gene-content-based phylogenetic reconstruction (Delsuc et al. 2005; Pett et al. 2019), we followed Zhao et al. (2021) to assign gene families for four species with OrthoFinder v2.4.0 (Emms and Kelly 2019) and then used the gene presence/absence matrix of 4 rows × 24,427 columns to construct a phylogenetic tree with IQ-TREE v2.1.2 (Minh et al. 2020) using the same model as the whole-genome microsynteny-based method.

The conventional coalescent approach based on single-copy orthologous genes was applied to reconstruct phylogenetic trees within 13 individuals of 13 species, *C. sinensis*, 6 EA *Carya* species (*C. cathayensis*, *C. dabieshanensis*, *C. hunanensis*, *C. kweichowensis*, *C. poilanei*, *C. tonkinensis*), 5 NA *Carya* species (*C. aquatica*, *C. cordiformis*, *C. illinoinensis, C. laciniosa*, *C. ovata*), and *P. stenoptera* (Supplementary Tables S4-S6)). First, the whole-genome protein sequences of four species (*C. sinensis*, *C. cathayensis*, *C. illinoinensis*, and *P. stenoptera*) were used with OrthoFinder v2.4.0 (Emms and Kelly 2019) to obtain 11,269 single-copy gene families. Second, we extracted the single-copy genes of the above-mentioned 13 species from the consensus genome, respectively. Third, each of the single-copy gene family protein sequences was aligned by MAFFT v7.273 (Katoh and Standley 2013) and then converted into corresponding codon alignments by PAL2NAL v14 (Suyama et al. 2006). The single-copy genes were filtered to satisfy the following cut-offs: sequence length between 300 and 2,000 bp and less than 20% gaps and 50% missing. PhiPack v1.1 (Bruen et al. 2006) was used to test for recombination for each single-copy gene family, using a permutation test with a *P*-value of <0.05 being treated as the recombinant gene. After filtering, a total of 7,398 single-copy genes from the consensus genome of 13 species (including *P. stenoptera*) were retained to perform the nuclear phylogenetic analysis. Gene trees were constructed by IQ-TREE v2.1.2 (Minh et al. 2020) using the maximum likelihood method with the parameters setting “-m MFP -B 1000” and the coalescent-based species tree was inferred by ASTRAL v5.7.4 (Mirarab et al. 2014).

#### *Introgression Test Based on the Species Tree of Carya sinensis and the 11 Diploid Carya Species*

Patterson's *D*-statistic (ABBA-BABA test) (Green et al. 2010; Durand et al. 2011) method was applied to the 30 rooted quartets of [((P1: *C. sinensis*, P2: EA *Carya* species), P3: NA *Carya* species), O: *P. stenoptera*], based on 11,269 single-copy gene families. The R package evobiR (Durand et al. 2011; Eaton and Ree 2013) was used, and the block (10-kb) bootstrap was applied to determine the significance of the *D*-statistic. The values obtained from the *D*-statistic test were all significantly different from zero (Supplementary Table S7), suggesting that gene flow could have occurred between EA *Carya* species (P2) and NA *Carya* species (P3), or from the ghost to *C. sinensis* (P1). To investigate the possibility of ghost introgression into *C. sinensi*s (P1), we employed the full-likelihood program BPP (Bayesian Phylogenetics and Phylogeography), which uses the multispecies-coalescent-with-introgression (MSci) model and assumes episodic introgression (Flouri et al. 2020). Specifically, we compared six introgression scenarios (Supplementary Fig. S6) for three triplets, [((*C. sinensis*, *C. cathayensis*), *C. illinoinensis*)], [((*C. sinensis*, *C. kweichowensis*), *C. cordiformis*)], [((*C. sinensis*, *C. tonkinensis*), *C. illinoinensis*)]. BPP assumes free recombination among loci and no recombination within a locus and, most remarkably, requires a lot of computation. Therefore, we only included those high-quality sequence alignments of single-copy gene families between 500 and 1,000 bp with less than 20% gaps and 50% missing. We then used PhiPack v1.1 (Bruen et al. 2006) to detect recombination for each single-copy gene family, using a permutation test with a P-value of <0.05 as the threshold for identifying recombinant genes. Ultimately, 2,543 single-copy gene families were used for BPP analysis, and we compared the six introgression models using marginal likelihood values calculated with BPP v4.6.2 (Flouri et al. 2020).

PhyloNet is used to infer species phylogenies while accounting for incomplete lineage sorting (ILS) and gene flow. We used PhyloNet v3.8.2 (Wen et al. 2018) to reconstruct phylogenetic networks from gene trees under a maximum pseudo-likelihood approach (Yu and Nakhleh 2015) with the command “InferNetwork_MPL”. We constructed 7,398 gene trees for each single-copy gene family using IQ-TREE. Network searches were performed using only nodes in the rooted ML gene trees with bootstrap support of at least 70% (-b 70), allowing for 0–3 reticulations in 25 runs (-X 25) and optimizing the branch lengths and inheritance probabilities (-o). Ten independent runs were carried out, and the run with the highest log probability was used.

#### *Divergence Time Estimates for Carya*

We estimated divergence times under the multispecies coalescent (MSC) in BPP based on a fixed species phylogeny inferred by ASTRAL for 12 species and the multispecies-coalescent-with-introgression (MSci) model in BPP based on the presence of ghost introgression from an extinct ancestor into the *C. sinensis*. To facilitate computation, we use a total of 773 non-recombinant single-copy gene families with sequence alignment lengths between 800 and 1,000 bp in this BPP analysis. Both analyses considered the heterozygote sites, using 500,000 MCMC iterations as burn-in and then taking 10,000 samples, sampling every 200 iterations in two different runs. The relative node ages (*θ*) estimated by BPP were rescaled from the substitution rate of 1.6×10^-9^ per site per year (Ding et al. 2023) to absolute time.

#### *Adaptive Introgression Genes and Gene Ontology Enrichment Analysis*

The genome-wide scans of adaptive introgression were implemented using VolcanoFinder v1.0 (Setter et al. 2020), which could be used to detect the signatures of ghost lineages based on the population genetic composite-likelihood method. Based on 24,751,236 SNPs from 43 individuals of *C. sinensis*, we generated the allele frequency file and the unnormalized site frequency spectrum using SweepFinder2 (DeGiorgio et al. 2016). The two files were used as inputs in the program VolcanoFinder with the options of “VolcanoFinder –ig 20000 and the Model 1” to identify the regions of adaptive introgression. The R package clusterProfiler (Yu et al. 2012) was used to perform gene ontology (GO) enrichment analysis for genes of possible adaptive introgression in the *C. sinensis* genome. The Benjamini-Hochberg method was used for multi-test correction, and subsequently, the overrepresented GO terms were chosen based on a false discovery rate (FDR) threshold of less than 0.05.

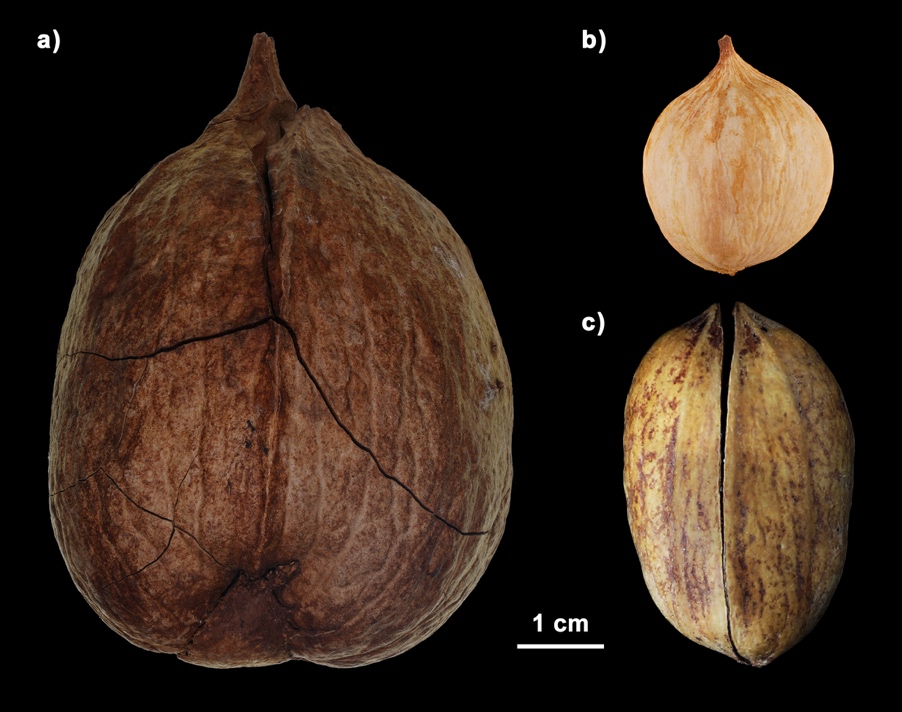

### Supplementary Figure 1. Shell morphology of *Carya sinensis* (a), *C. cathayensis* (b), and *C. illinoinensis* (c).

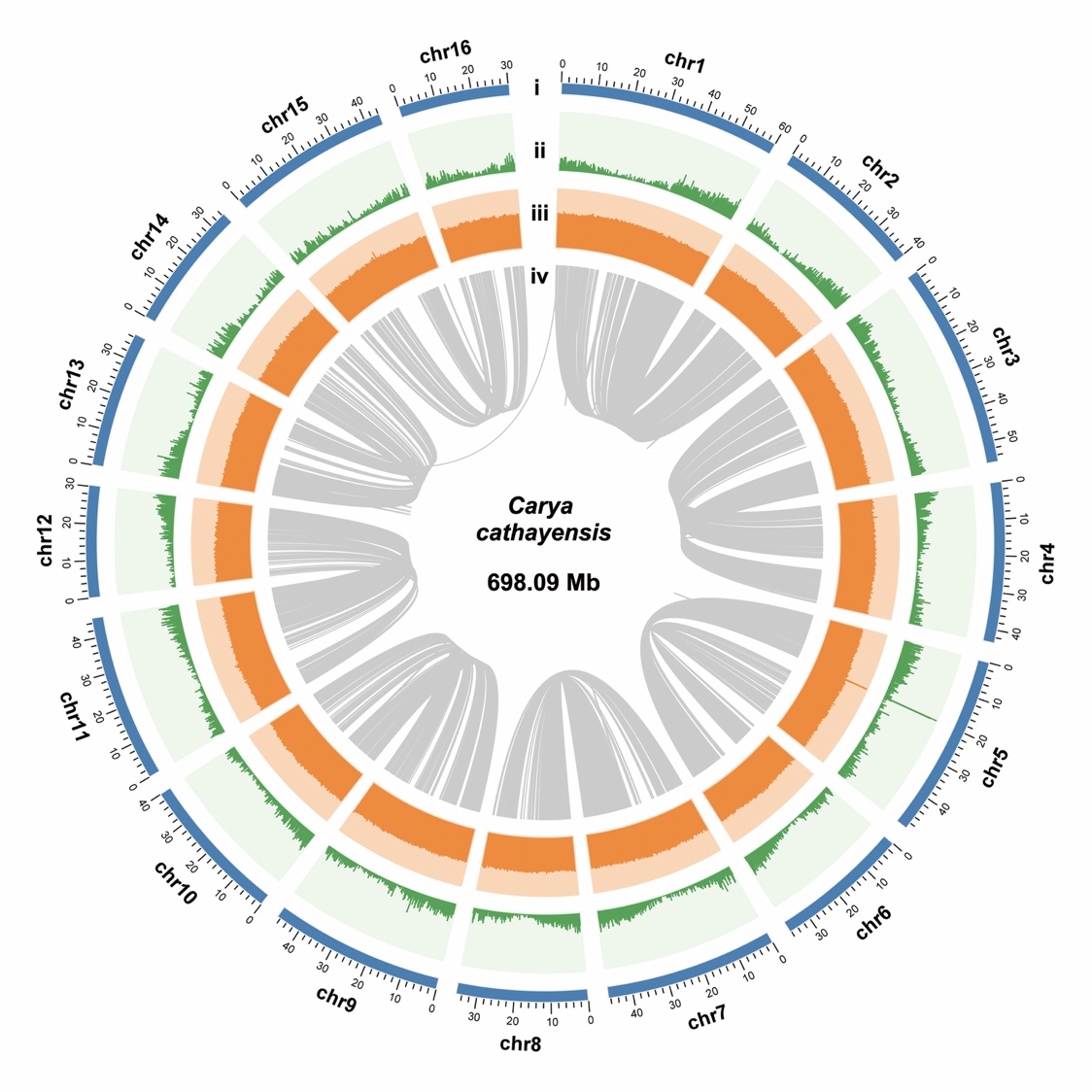

### Supplementary Figure 2. Genome features of *Carya cathayensis.* Different tracks (moving inward) denote (i) 16 chromosomes; (ii) gene density in 100 kb stepping windows (minimum-maximum, 1–47); (iii) GC content in 100 bp stepping windows (minimum-maximum, 0–0.64); (iv) identified syntenic blocks.

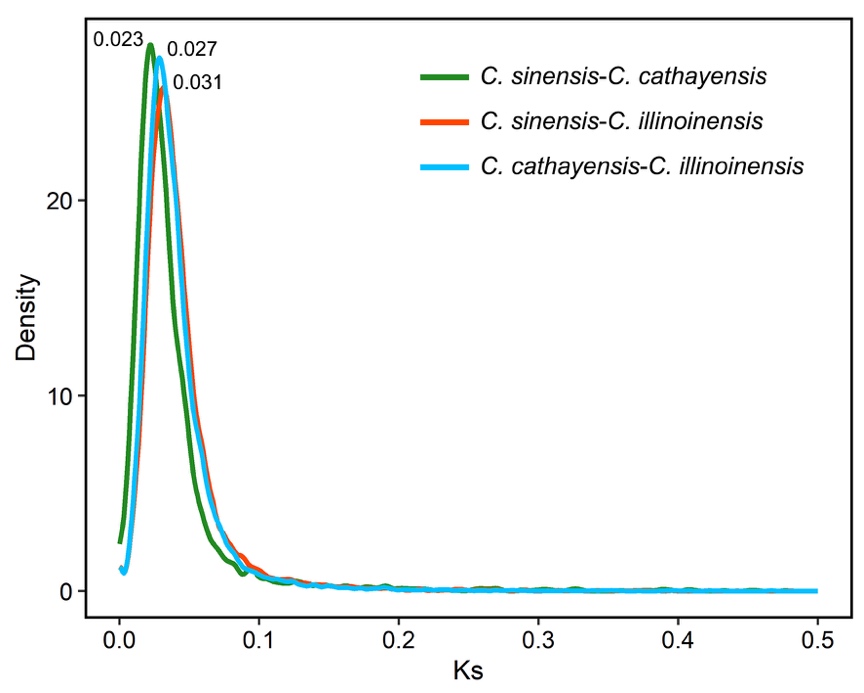

### Supplementary Figure 3. Synonymous substitution rate (*K*_s_) density distributions of syntenic orthologs among *Carya sinensis*, *C. cathayensis* and *C. illinoinensis*.

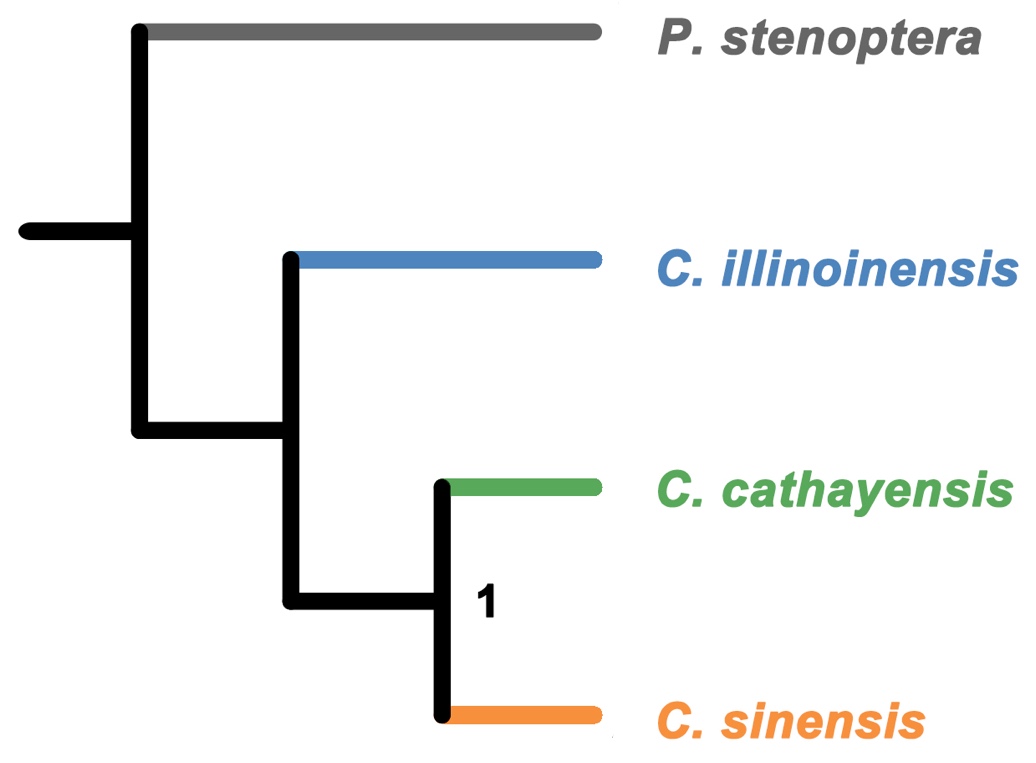

### Supplementary Figure 4. Species tree reconstructed for *Pterocarya stenoptera*, *Carya sinensis,* *C. cathayensis*, and *C. illinoinensis* using three methods of DNA-sequence alignment with ASTRAL-Pro, whole-genome microsynteny, and local gene content.

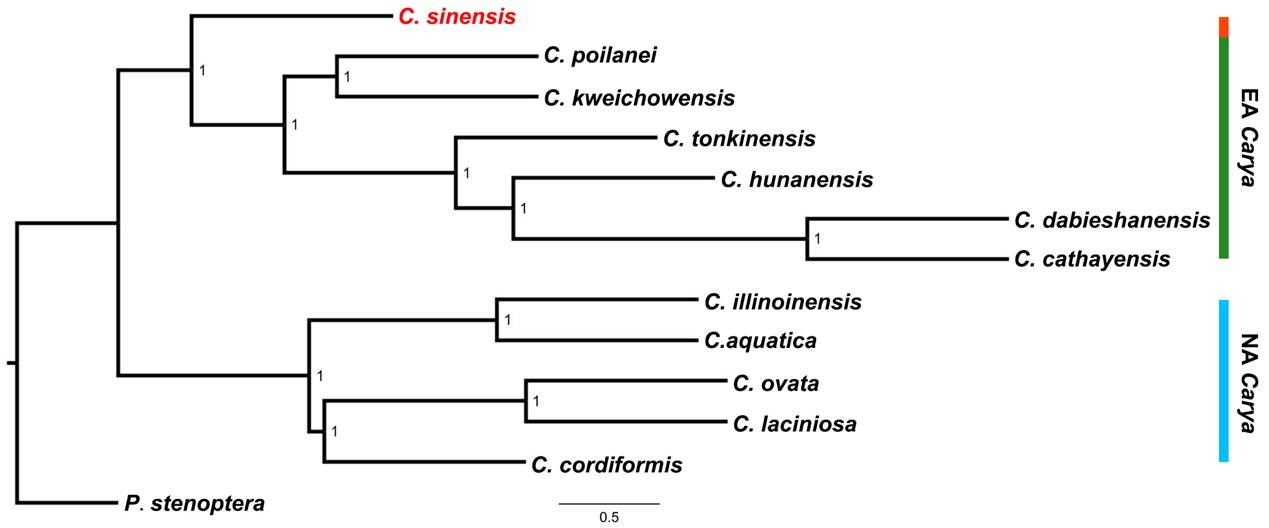

### Supplementary Figure 5. Species tree of *Carya sinensis* and 11 diploid *Carya* species with *Pterocarya stenoptera* as outgroup inferred using ASTRAL based on 7,398 single-copy nuclear genes.

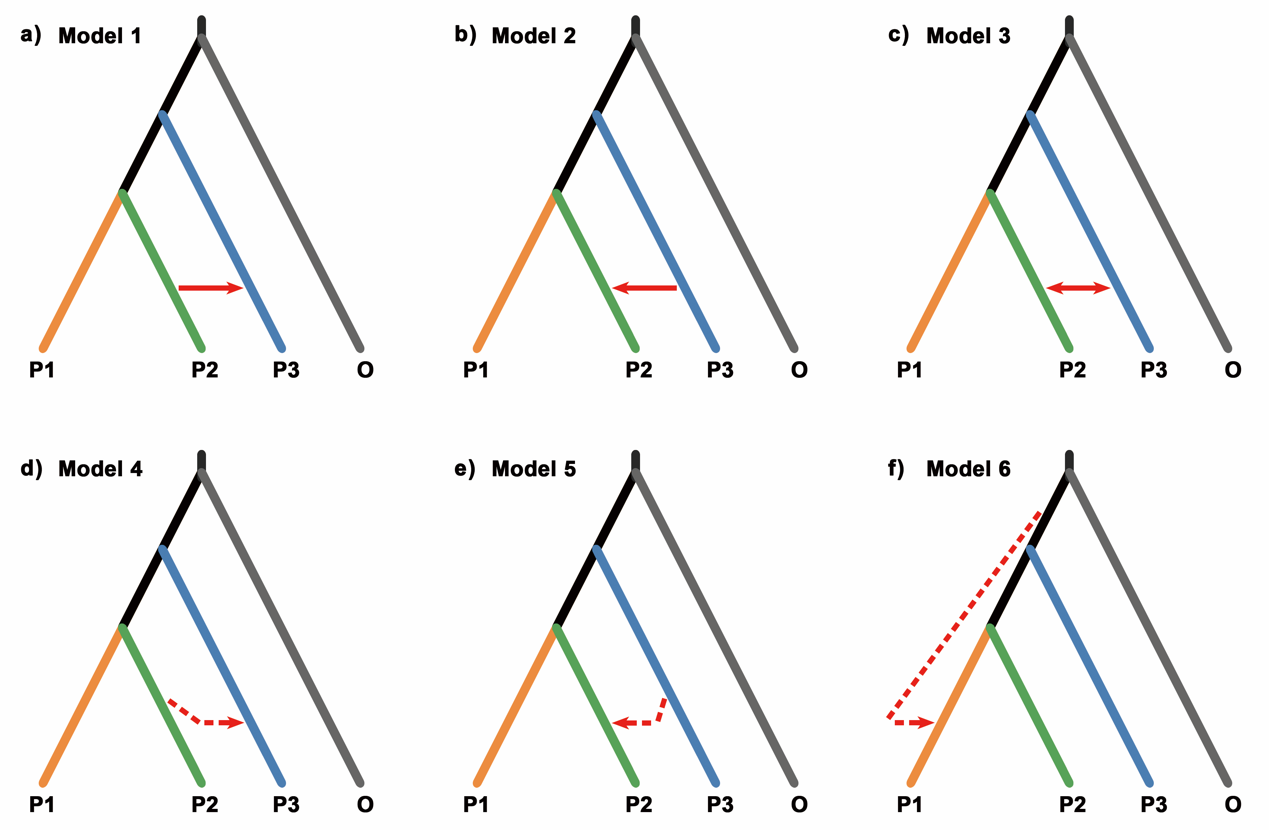

### Supplementary Figure 6. Six introgression scenarios based on a ladder-like species tree of four extant lineages (ingroups: P1, P2, and P3 and outgroup: O) and one ghost lineage. The red solid arrows indicate the introgression events between the extant lineage P2 and P3 (a-c); and the red dotted arrows indicate the ghost introgression events from ingroup ghost lineage to P3 (e), P2 (f) or outgroup ghost lineage to P1 (g), respectively. The six introgression events (a-f) can result in a significant positive *D*-statistic.

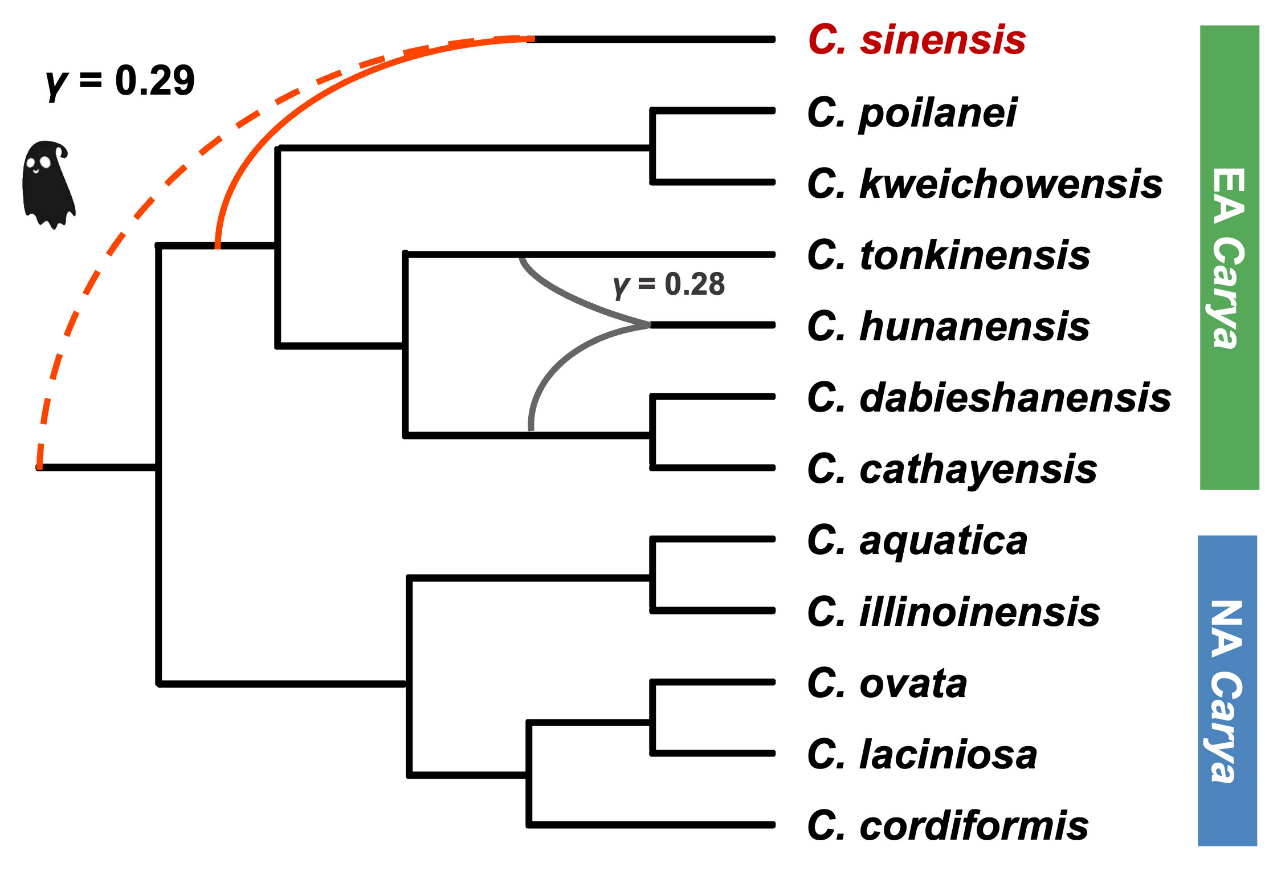

### Supplementary Figure 7. Species network inference of *Carya sinensis* and 11 diploid species of genus *Carya.* The result was inferred using PhyloNet based on 7,398 single-copy genes with two reticulations. Grey and red curves represent the hybridization events of *C. hunanensis* (one reticulation) and *C. sinensis* (two reticulations), respectively. The γ value indicates the inheritance probability from donor lineage to recipient lineage.

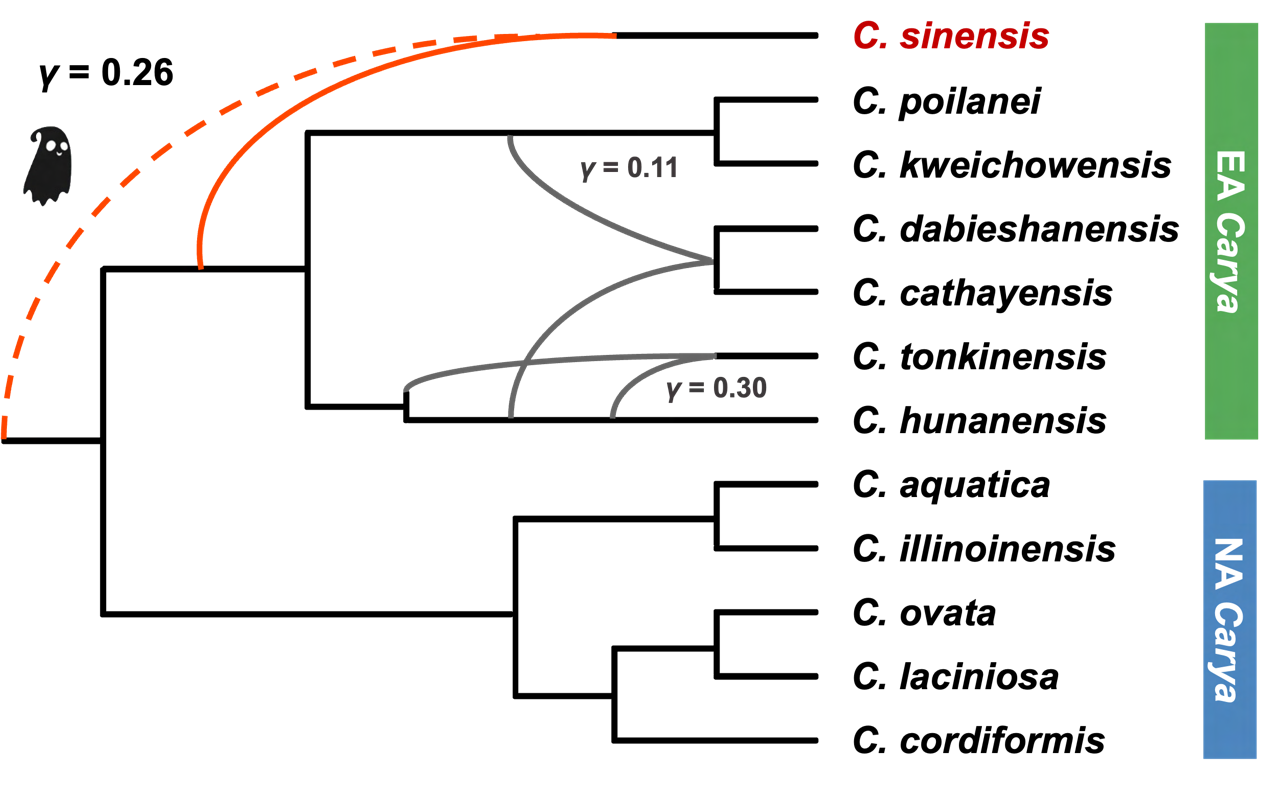

### Supplementary Figure 8. Species network with three reticulations of *Carya sinensis* and 11 diploid *Carya* species inferred using PhyloNet-MPL based on 7,398 single-copy nuclear genes. The *γ* values indicate the inheritance probability from the parental species, the red curve represents the hybridization events of *C. sinensis*, and the grey curves represent the hybridization events of other EA *Carya* species.

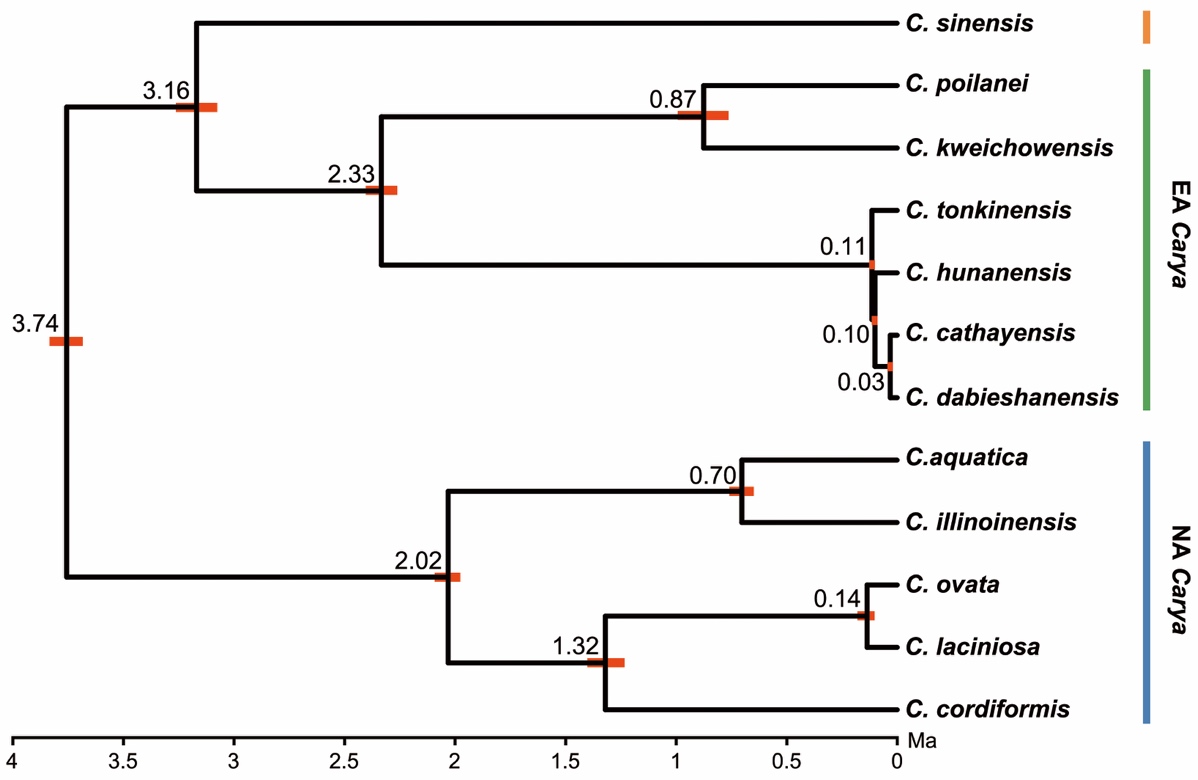

### Supplementary Figure 9. The estimated divergence times under the multispecies coalescent (MSC) in BPP are based on the *Carya sinensis* and the other 11 diploid *Carya* species. Node heights are posterior means under the MSC model with the absolute divergence times. Posterior means and 95% HPD CIs (red bars) are shown above their vertices to improve visibility.

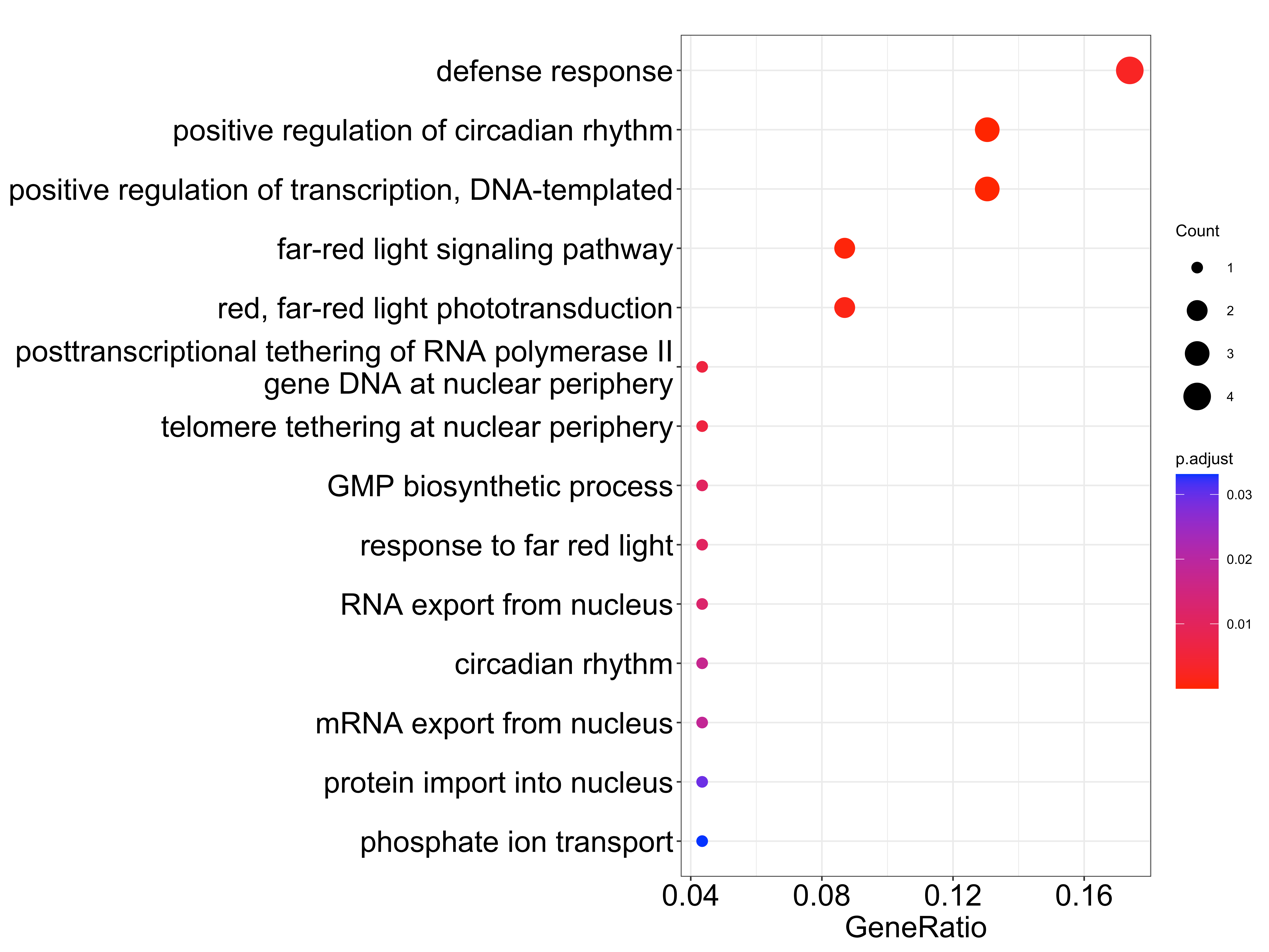

### Supplementary Figure 10. Gene ontology enrichment analysis of 36 possible ghost introgression genes detected in the *Carya sinensis* genome.

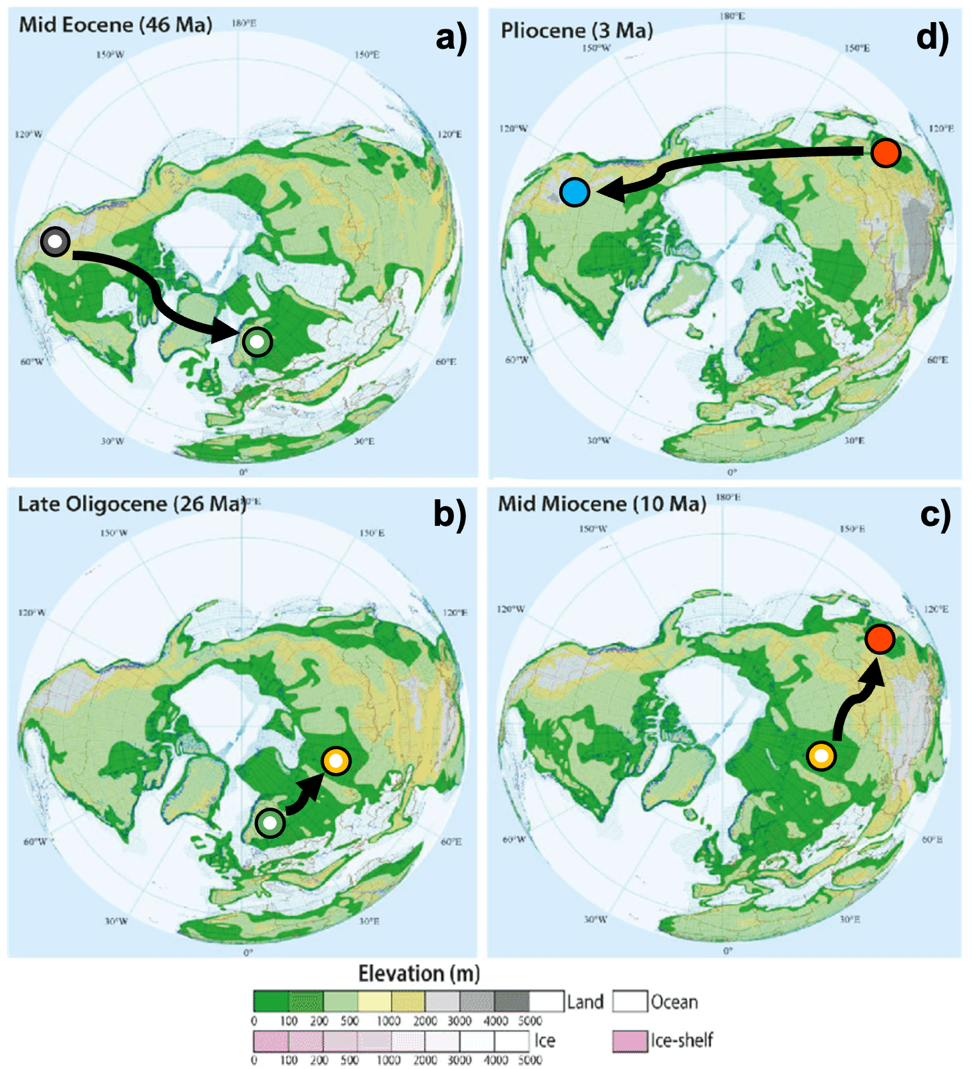

### Supplementary Figure 11. Modified hypothetical migratory routes of *Carya* in the biogeographic scenario. The geographical pattern of the Beringian Land Bridge (BLB) in different periods was from Wen et al. (2016). The migratory routes of *Carya* in Mid Eocene(a), Late Oligocene (b), and Mid Miocene (c) were adapted from Zhang et al. (2013), and in Pliocene (d) was a new one we added. The hollow and solid circles indicated the representative geographic distribution areas of ancient fossils and living species of *Carya*, respectively.

### Supplementary Table 1. Assembly and annotation features of two pseudo-chromosomal level genomes.

| **Genome Statistics** | ***Carya sinensis*** | ***Carya cathayensis*** |
| --- | --- | --- |
| Illumina short reads (Gb) | 33.88 | 36.66 |
| PacBio single-molecule long reads (Gb) | 49.51 | 63.19 |
| Hi-C interaction reads (Gb) | 152.12 | 107.68 |
| Estimated genome size (Mb) | 634.52 | 742.32 |
| Assembled genome size (Mb) | 623.16 | 698.09 |
| No. of scaffolds (>10kb) | 226 | 198 |
| *N*_50_ length (scaffold) (Mb) | 38.85 | 43.49 |
| *L*_50_ (scaffold) | 8 | 8 |
| Longest scaffold (Mb) | 54.83 | 60.01 |
| Chromosome anchor percent (%) | 96.5 | 95.9 |
| GC content (%) | 0.36 | 0.36 |
| Transposable elements (%) | 53.30 | 58.91 |
| Predicted protein-coding genes | 35,370 | 36,722 |

### Supplementary Table 2. Genome completeness of *Carya sinensis* and *C. cathayensis* was measured by Benchmarking Universal Single-Copy Orthologs (BUSCO) within 1,440 single copy ortholog groups in plants.

| **BUSCO notation assessment** | ***Carya sinensis*** | ***Carya cathayensis*** |
| --- | --- | --- |
| **Complete BUSCOs** | 94.5% | 93.9% |
| **Complete and Single-copy BUSCOs** | 83.0% | 80.4% |
| **Complete and Duplicated BUSCOs** | 11.5% | 13.5% |
| **Fragmented BUSCOs** | 1.3% | 1.8% |
| **Missing BUSCOs** | 4.2% | 4.3% |

### Supplementary Table 3. Functional annotation of the predicted genes models.

| **Type** | ***Carya sinensis*** | | | ***Carya cathayensis*** | |
| --- | --- | --- | --- | --- | --- |
|  | **Number** | **Percent (%)** | **Number** | | **Percent (%)** |
| **GO** | 23,790 | 67.26% | 24,477 | | 66.65% |
| **KEGG** | 31,654 | 89.49% | 32,323 | | 88.02% |
| **NCBI-NR** | 31,609 | 89.37% | 32,248 | | 87.82% |
| **UniProt-TrEMBL** | 31,831 | 89.99% | 32,444 | | 88.35% |
| **Total** | **31,951** | **90.33%** | **32,593** | | **88.76%** |

### Supplementary Table 4. The re-sequencing data of 43 *Carya sinensis* individuals included in this study, with sample id, location, and coverage rate of its reference genome.

| **No.** | **Sample ID** | **Herbarium voucher** | **Latitude (N)** | **Longitude (E)** | **Location** | **Depth (×)** | **Coverage (>1×)** | **Coverage (1/3** – **2× of depth)** |
| --- | --- | --- | --- | --- | --- | --- | --- | --- |
| 1 | Asi-AL1 | 20190730-18 | 25.16953 | 105.64366 | Anlong, Guizhou, China | 29.61 | 0.961 | 0.901 |
| 2 | Asi-CH1 | GZBG02201 | 24.92478 | 105.84342 | Ceheng, Guizhou, China | 26.02 | 0.96 | 0.901 |
| 3 | Asi-FN1 | 20180709-1 | 23.58113 | 105.5808 | Funing, Yunnan, China | 26.86 | 0.96 | 0.904 |
| 4 | Asi-FN2 | 20180709-2 | 23.58113 | 105.5808 | Funing, Yunnan, China | 29.43 | 0.95 | 0.889 |
| 5 | Asi-HK1 | ZJULP03501 | 22.67474 | 104.02768 | Hekou, Yunnan, China | 30.98 | 0.962 | 0.897 |
| 6 | Asi-HK11 | ZJULP03712 | 22.69106 | 104.02379 | Hekou, Yunnan, China | 26.58 | 0.963 | 0.903 |
| 7 | Asi-HK13 | ZJULP03715 | 22.69106 | 104.02379 | Hekou, Yunnan, China | 30.61 | 0.963 | 0.902 |
| 8 | Asi-HK15 | ZJULP03803 | 22.66436 | 104.01284 | Hekou, Yunnan, China | 26.11 | 0.96 | 0.902 |
| 9 | Asi-LB1 | 20180718-1 | 25.22007 | 107.97092 | Libo, Guizhou, China | 24.12 | 0.959 | 0.889 |
| 10 | Asi-LB10 | GZBG03401 | 25.27705 | 107.92471 | Libo, Guizhou, China | 24.17 | 0.958 | 0.89 |
| 11 | Asi-LB2 | 20180718-2 | 25.21524 | 107.98218 | Libo, Guizhou, China | 20.7 | 0.957 | 0.895 |
| 12 | Asi-LB3 | 20180719-17 | 25.25429 | 107.72696 | Libo, Guizhou, China | 22.51 | 0.956 | 0.896 |
| 13 | Asi-LB6 | 20180719-20 | 25.25429 | 107.72696 | Libo, Guizhou, China | 20.8 | 0.957 | 0.898 |
| 14 | Asi-LB7 | 20180719-21 | 25.25429 | 107.72696 | Libo, Guizhou, China | 21.08 | 0.956 | 0.887 |
| 15 | Asi-LB8 | GZBG03801 | 25.13603 | 107.82411 | Libo, Guizhou, China | 27.97 | 0.958 | 0.895 |
| 16 | Asi-LB9 | GZBG03802 | 25.13603 | 107.82411 | Libo, Guizhou, China | 23.61 | 0.956 | 0.896 |
| 17 | Asi-LC10 | 20180725-10 | 24.90306 | 108.57343 | Luocheng, Guangxi, China | 31.78 | 0.952 | 0.886 |
| 18 | Asi-LC3 | 20180725-3 | 24.90306 | 108.57343 | Luocheng, Guangxi, China | 25.97 | 0.961 | 0.897 |
| 19 | Asi-LD1 | GZBG04101 | 25.44326 | 106.48932 | Luodian, Guizhou, China | 24.19 | 0.959 | 0.894 |
| 20 | Asi-LD2 | GZBG04201 | 25.5295 | 106.44975 | Luodian, Guizhou, China | 25.91 | 0.959 | 0.898 |
| 21 | Asi-LY1 | 20200822-4 | 24.74204 | 106.5882 | Leyue, Guangxi, China | 27.46 | 0.959 | 0.891 |
| 22 | Asi-MLP1 | 20180708-1 | 23.13038 | 104.8585 | Malipo, Yunnan, China | 26.89 | 0.995 | 0.967 |
| 23 | Asi-MLP11 | 20180708-19 | 23.13038 | 104.8585 | Malipo, Yunnan, China | 25.9 | 0.96 | 0.902 |
| 24 | Asi-MLP12 | 20191116-1 | 23.00419 | 104.82261 | Malipo, Yunnan, China | 33.4 | 0.959 | 0.893 |
| 25 | Asi-MLP14 | 20191116-3 | 23.00419 | 104.82261 | Malipo, Yunnan, China | 31.8 | 0.966 | 0.909 |
| 26 | Asi-MLP15 | 20191116-4 | 23.00419 | 104.82261 | Malipo, Yunnan, China | 29.81 | 0.961 | 0.904 |
| 27 | Asi-MLP16 | 20191116-5 | 23.00419 | 104.82261 | Malipo, Yunnan, China | 27.22 | 0.96 | 0.897 |
| 28 | Asi-MLP17 | ZJUQXY00701 | 23.18185 | 104.82005 | Malipo, Yunnan, China | 27.78 | 0.963 | 0.901 |
| 29 | Asi-MLP4 | 20180708-4 | 23.13038 | 104.8585 | Malipo, Yunnan, China | 26.78 | 0.96 | 0.9 |
| 30 | Asi-NP1 | 20190726-11 | 23.50354 | 105.83882 | Napo, Guangxi, China | 33.54 | 0.964 | 0.891 |
| 31 | Asi-NP2 | 20190726-12 | 23.50354 | 105.83882 | Napo, Guangxi, China | 30 | 0.962 | 0.9 |
| 32 | Asi-NP3 | 20190726-13 | 23.50354 | 105.83882 | Napo, Guangxi, China | 24.23 | 0.962 | 0.895 |
| 33 | Asi-RJ1 | GZBG01201 | 25.88452 | 108.55526 | Rongjiang, Guizhou, China | 24.76 | 0.958 | 0.893 |
| 34 | Asi-SD15 | 20180720-16 | 26.03958 | 107.97767 | Sandu, Guizhou, China | 27.01 | 0.952 | 0.882 |
| 35 | Asi-SD2 | 20180720-2 | 26.03958 | 107.97767 | Sandu, Guizhou, China | 19.15 | 0.957 | 0.889 |
| 36 | Asi-SD3 | 20180720-3 | 26.03958 | 107.97767 | Sandu, Guizhou, China | 26.84 | 0.959 | 0.901 |
| 37 | Asi-TD1 | 20180722-6 | 25.89419 | 109.74927 | Tongdao, Hunan, China | 17.14 | 0.952 | 0.871 |
| 38 | Asi-WM1 | 20180714-1 | 25.15908 | 106.20026 | Wangmo, Guizhou, China | 20.42 | 0.955 | 0.893 |
| 39 | Asi-XC1 | 20190423-3 | 23.43976 | 104.85461 | Xichou, Yunnan, China | 28.32 | 0.965 | 0.902 |
| 40 | Asi-XC2 | ZJUQXY01201 | 23.44728 | 104.82088 | Xichou, Yunnan, China | 25.17 | 0.96 | 0.9 |
| 41 | Asi-XC4 | ZJUQXY01302 | 23.44577 | 104.82948 | Xichou, Yunnan, China | 27.61 | 0.962 | 0.898 |
| 42 | Asi-XC6 | ZJUQXY01401 | 23.46901 | 104.81338 | Xichou, Yunnan, China | 27.96 | 0.958 | 0.895 |
| 43 | Asi-XC7 | ZJUQXY01501 | 23.46207 | 104.8621 | Xichou, Yunnan, China | 28.68 | 0.96 | 0.9 |

### Supplementary Table 5. The re-sequencing data of 6 East Asia *Carya* individuals included in this study, with sample id, location, and coverage rate of *C. cathayensis* reference genome.

| **No.** | **Sample ID** | **Species** | **Herbarium voucher** | **Latitude (N)** | **Longitude (E)** | **Location** | **Depth (×)** | **Coverage (>1×)** | **Coverage (1/3** – **2× of depth)** |
| --- | --- | --- | --- | --- | --- | --- | --- | --- | --- |
| 1 | Cca-QLF12 | *C. cathayensis* | 20180829-47 | 30.14208 | 118.84434 | Jixi, Anhui, China | 24.55 | 0.997 | 0.957 |
| 2 | Cda-JZ4 | *C. dabieshanensis* | 20190721-107 | 31.25944 | 115.83923 | Jinzhai, Anhui, China | 30.85 | 0.964 | 0.892 |
| 3 | Chu-TD2 | *C. hunanensis* | 20190805-112 | 26.11490 | 109.69434 | Tongdao, hunan, China | 26.71 | 0.966 | 0.850 |
| 4 | Ckw-CH1 | *C. kweichowensis* | 20200824-1 | 25.07081 | 105.81433 | Ceheng, Guizhou, China | 24.62 | 0.802 | 0.569 |
| 5 | Cpo-XMT1 | *C. poilanei* | 20210731-1 | 23.40167 | 102.87152 | Jianshui, Yunnan, China | 24.83 | 0.758 | 0.556 |
| 6 | Cto-YP1 | *C. tonkinensis* | 20190825-1 | 25.13847 | 99.54054 | Yongping, Yunnan, China | 26.74 | 0.832 | 0.648 |

### Supplementary Table 6. The re-sequencing data of 5 North America *Carya* individuals included in this study, with sample id, location, ploidy level and coverage rate of *C. illinoinensis* reference genome.

| **No.** | **Sample ID** | **Species** | **Herbarium voucher** | **Location** | **Depth (×)** | **Coverage (>1×)** | **Coverage (1/3** – **2× of depth)** | **Ploidy**  **level** |
| --- | --- | --- | --- | --- | --- | --- | --- | --- |
| 1 | Caq-DA1 | *C. aquatica* | Caq-14368 | AL, Dallas Co, USA | 27.73 | 0.898 | 0.721 | diploid |
| 2 | Cco-WI3 | *C. cordiformis* | Cco-13122 | AL, Wilcox Co., USA | 28.25 | 0.816 | 0.597 | diploid |
| 3 | Cil-MA2 | *C. illinoinensis* | LS20191026-01 | United States of America, Iowa, Des Moines, on the Mississippi River bottom near Burlington | 23.38 | 0.957 | 0.854 | diploid |
| 4 | Cla-2884 | *C. laciniosa* | Cla-2884 | cultivate in Munich Botanical Garden | 27.93 | 0.808 | 0.577 | diploid |
| 5 | Cov-MA1 | *C. ovata* | LS20191025-04 | United States of America, Illinois, Macon, 0.25 mile south of the NE corner of Spitler Woods State Park on the east edge of the park | 24.28 | 0.803 | 0.580 | diploid |

### Supplementary Table 7. Results from Patterson’s *D* test for introgression between species with SD estimates and significance values.

| **No.** | **P1** | **P2** | **P3** | ***D*-statistic** | **block bootstrap SD** | **Z-score** | **P-value** |
| --- | --- | --- | --- | --- | --- | --- | --- |
| 1 | *C. sinensis* | *C. poilanei* | *C. aquatica* | 0.0966449 | 0.008264317 | 11.69424 | 0 |
| 2 | *C. sinensis* | *C. poilanei* | *C. illinoinensis* | 0.09784537 | 0.008032938 | 12.18052 | 0 |
| 3 | *C. sinensis* | *C. poilanei* | *C. ovata* | 0.1052412 | 0.008170307 | 12.88094 | 0 |
| 4 | *C. sinensis* | *C. poilanei* | *C. laciniosa* | 0.1044981 | 0.008189018 | 12.76076 | 0 |
| 5 | *C. sinensis* | *C. poilanei* | *C. cordiformis* | 0.09545638 | 0.008305279 | 11.49346 | 0 |
| 6 | *C. sinensis* | *C. kweichowensis* | *C. aquatica* | 0.09216795 | 0.008174147 | 11.27554 | 0 |
| 7 | *C. sinensis* | *C. kweichowensis* | *C. illinoinensis* | 0.09916532 | 0.007958934 | 12.45962 | 0 |
| 8 | *C. sinensis* | *C. kweichowensis* | *C. ovata* | 0.1015662 | 0.007958307 | 12.76229 | 0 |
| 9 | *C. sinensis* | *C. kweichowensis* | *C. laciniosa* | 0.1026472 | 0.008513823 | 12.05654 | 0 |
| 10 | *C. sinensis* | *C. kweichowensis* | *C. cordiformis* | 0.09420194 | 0.008463396 | 11.13051 | 0 |
| 11 | *C. sinensis* | *C. tonkinensis* | *C. aquatica* | 0.07703963 | 0.007551766 | 10.20154 | 0 |
| 12 | *C. sinensis* | *C. tonkinensis* | *C. illinoinensis* | 0.07899851 | 0.00773225 | 10.21676 | 0 |
| 13 | *C. sinensis* | *C. tonkinensis* | *C. ovata* | 0.07666185 | 0.007587375 | 10.10387 | 0 |
| 14 | *C. sinensis* | *C. tonkinensis* | *C. laciniosa* | 0.07465095 | 0.007884952 | 9.46752 | 0 |
| 15 | *C. sinensis* | *C. tonkinensis* | *C. cordiformis* | 0.07404245 | 0.007692994 | 9.62466 | 0 |
| 16 | *C. sinensis* | *C. hunanensis* | *C. aquatica* | 0.1455108 | 0.008229166 | 17.68233 | 0 |
| 17 | *C. sinensis* | *C. hunanensis* | *C. illinoinensis* | 0.1474538 | 0.008075534 | 18.25932 | 0 |
| 18 | *C. sinensis* | *C. hunanensis* | *C. ovata* | 0.1681465 | 0.008460379 | 19.87458 | 0 |
| 19 | *C. sinensis* | *C. hunanensis* | *C. laciniosa* | 0.1640449 | 0.008198097 | 20.01012 | 0 |
| 20 | *C. sinensis* | *C. hunanensis* | *C. cordiformis* | 0.1562121 | 0.008504121 | 18.36898 | 0 |
| 21 | *C. sinensis* | *C. dabieshanensis* | *C. aquatica* | 0.09264842 | 0.007853696 | 11.79679 | 0 |
| 22 | *C. sinensis* | *C. dabieshanensis* | *C. illinoinensis* | 0.09465067 | 0.007487807 | 12.64064 | 0 |
| 23 | *C. sinensis* | *C. dabieshanensis* | *C. ovata* | 0.09983776 | 0.007829086 | 12.75216 | 0 |
| 24 | *C. sinensis* | *C. dabieshanensis* | *C. laciniosa* | 0.09684172 | 0.007680103 | 12.60943 | 0 |
| 25 | *C. sinensis* | *C. dabieshanensis* | *C. cordiformis* | 0.09471259 | 0.007917236 | 11.96284 | 0 |
| 26 | *C. sinensis* | *C. cathayensis* | *C. aquatica* | 0.08904386 | 0.007380431 | 12.06486 | 0 |
| 27 | *C. sinensis* | *C. cathayensis* | *C. illinoinensis* | 0.09107334 | 0.007575985 | 12.02132 | 0 |
| 28 | *C. sinensis* | *C. cathayensis* | *C. ovata* | 0.09490225 | 0.007697483 | 12.329 | 0 |
| 29 | *C. sinensis* | *C. cathayensis* | *C. laciniosa* | 0.09278602 | 0.007419295 | 12.50604 | 0 |
| 30 | *C. sinensis* | *C. cathayensis* | *C. cordiformis* | 0.09108411 | 0.007737844 | 11.77125 | 0 |

### Supplementary Table 8. Marginal log-likelihood values in BPP analyses of the six introgression scenarios.

| **Triplets** | **log marginal likelihood** | | | | | |
| --- | --- | --- | --- | --- | --- | --- |
|  | **Model 1** | **Model 2** | **Model 3** | **Model 4** | **Model 5** | **Model 6** |
| ((*C. sinensis*, *C. cathayensis*), *C. illinoinensis*) | -3,138,493 | -3,138,589 | -3,138,489 | -3,138,468 | -3,138,512 | -3,138,311 |
| ((*C. sinensis*, *C. kweichowensis*), *C. cordiformis*) | -3,096,025 | -3,096,261 | -3,096,095 | -3,096,058 | -3,096,237 | -3,095,860 |
| ((*C. sinensis*, *C. tonkinensis*), *C. illinoinensis*) | -3,139,990 | -3,140,206 | -3,140,206 | -3,140,079 | -3,140,154 | -3,139,789 |

### Supplementary Table 9. Possible ghost introgression genes detected by VolcanoFinder in *Carya sinensis* genome.

| **Chromosome** | **Position** | **CLR** | **Gene** |
| --- | --- | --- | --- |
| Chr14 | 6089347 | 302.7351 | ASI011833 |
| Chr08 | 14550404 | 238.9519 | ASI034699 |
| Chr14 | 15057837 | 231.003 | ASI034252 |
| Chr14 | 15037818 | 218.2205 | NA |
| Chr14 | 6109366 | 217.2463 | NA |
| Chr03 | 11348878 | 213.3127 | ASI034477, ASI034478, ASI034480 |
| Chr08 | 14530390 | 210.7706 | NA |
| Chr14 | 15077856 | 206.6819 | ASI034252 |
| Chr11 | 9948655 | 196.0625 | ASI031967 |
| Chr14 | 15017799 | 195.8947 | NA |
| Chr14 | 15097875 | 189.6494 | NA |
| Chr09 | 16946219 | 184.4843 | ASI007222 |
| Chr14 | 14997780 | 169.7005 | NA |
| Chr01 | 30020100 | 168.5856 | NA |
| Chr09 | 16966226 | 151.6349 | NA |
| Chr10 | 13877118 | 148.311 | ASI012440 |
| Chr16 | 21799237 | 148.0938 | ASI024295 |
| Chr06 | 15092573 | 147.9993 | ASI014024 |
| Chr04 | 20373161 | 147.3524 | NA |
| Chr14 | 15117894 | 139.2846 | NA |
| Chr14 | 6069328 | 138.3498 | ASI011726, ASI011809, ASI011983 |
| Chr10 | 13897133 | 134.557 | NA |
| Chr15 | 26240658 | 130.7961 | ASI017001, ASI017037 |
| Chr03 | 11368890 | 130.3059 | NA |
| Chr15 | 18614862 | 127.8254 | ASI031428 |
| Chr15 | 29302986 | 127.3806 | ASI030466, ASI030485 |
| Chr15 | 29282971 | 126.0448 | ASI030466 |
| Chr15 | 37669344 | 124.8139 | ASI018686 |
| Chr15 | 29323001 | 124.6136 | NA |
| Chr15 | 29262955 | 122.9064 | ASI030479 |
| Chr15 | 29242940 | 119.282 | ASI030476, ASI030500, ASI030502 |
| Chr03 | 11589020 | 116.6026 | NA |
| Chr08 | 14570418 | 115.8047 | NA |
| Chr10 | 26806879 | 114.7815 | NA |
| Chr15 | 18594847 | 114.5424 | ASI031436, ASI031447, ASI031453 |
| Chr15 | 29343016 | 114.13 | ASI030454, ASI030493 |
| Chr15 | 29222925 | 113.7952 | ASI030450, ASI030503 |
| Chr12 | 3102266 | 112.4618 | NA |
| Chr15 | 26260673 | 111.5238 | ASI016994 |
| Chr12 | 3082253 | 111.238 | ASI026022 |
| Chr15 | 29202910 | 108.8282 | ASI030459, ASI030482, ASI030503 |
| Chr16 | 25102619 | 107.8569 | ASI033493 |
| Chr10 | 13476815 | 105.319 | NA |
| Chr15 | 29182894 | 101.2928 | ASI030459 |

### Supplementary Table 10. The Gene function of possible ghost introgression genes detected by VolcanoFinder in *Carya sinensis* genome.

| **Gene** | **GO Term** | **Gene function (against the KEGG databases)** |
| --- | --- | --- |
| ASI011809.1 | GO:0016021 | uncharacterized LOC101304047 |
| ASI011833.1 | GO:0005509 | calcineurin B-like protein 1 isoform X1 |
| ASI011983.1 | GO:0006952; GO:0043531 | putative disease resistance protein RGA3 |
| ASI014024.1 | GO:0016021 | UPF0481 protein At3g47200 |
| ASI016994.1 | GO:0005783; GO:0005886; GO:0006817; GO:0006888; GO:0016021 | SEC12-like protein 1 |
| ASI018686.1 | GO:0004970; GO:0016021 | glutamate receptor 2.7 |
| ASI024295.1 | GO:0006952; GO:0043531 | putative disease resistance RPP13-like protein 1 |
| ASI026022.1 | GO:0016491 | (-)-isopiperitenol/(-)-carveol dehydrogenase, mitochondrial-like |
| ASI030450.1 | GO:0016021 | UPF0481 protein At3g47200 |
| ASI030454.1 | GO:0016021 | UPF0481 protein At3g47200 |
| ASI030459.1 | GO:0016021 | transmembrane 9 superfamily member 8 |
| ASI030479.1 | GO:0016021 | UPF0481 protein At3g47200-like |
| ASI030485.1 | GO:0016021 | UPF0481 protein At3g47200 |
| ASI030493.1 | GO:0030001; GO:0046872 | heavy metal-associated isoprenylated plant protein 26 |
| ASI030500.1 | GO:0016021 | UPF0481 protein At3g47200 |
| ASI031447.1 | GO:0000973; GO:0003723; GO:0005487; GO:0006405; GO:0006406; GO:0006606; GO:0008139; GO:0009733; GO:0009870; GO:0017056; GO:0031965; GO:0034398; GO:0044614 | nuclear pore complex protein NUP96 |
| ASI031967.1 | GO:0000166; GO:0003938; GO:0005737; GO:0006177; GO:0046872 | inosine-5'-monophosphate dehydrogenase 2-like |
| ASI033493.1 | GO:0006952; GO:0007165; GO:0043531 | TMV resistance protein N |
| ASI034252.1 | GO:0006952; GO:0043531 | putative disease resistance gene NBS-LRR family protein |
| ASI034477.1 | GO:0003700; GO:0005634; GO:0008270; GO:0010018; GO:0042753; GO:0045893 | protein FAR-RED IMPAIRED RESPONSE 1 |
| ASI034478.1 | GO:0003700; GO:0005634; GO:0007623; GO:0008270; GO:0009585; GO:0010218; GO:0042753; GO:0045893 | protein FAR-RED ELONGATED HYPOCOTYL 3 |
| ASI034480.1 | GO:0003700; GO:0005634; GO:0008270; GO:0009585; GO:0010018; GO:0042753; GO:0045893 | protein FAR1-RELATED SEQUENCE 2 |
| ASI034699.1 | GO:0008270 | pentatricopeptide repeat-containing protein At3g63370, chloroplastic |

The grey highlighted table frames represent the genes that are associated with transcription factors FHY3 and FAR1*.*
